# A model for the emergence of RNA from a prebiotically plausible mixture of ribonucleotides, arabinonucleotides and 2’-deoxynucleotides

**DOI:** 10.1101/813675

**Authors:** Seohyun Chris Kim, Lijun Zhou, Wen Zhang, Derek K. O’Flaherty, Valeria Rondo-Brovetto, Jack W. Szostak

## Abstract

The abiotic synthesis of ribonucleotides is thought to have been an essential step towards the emergence of the RNA world. However, it is likely that the prebiotic synthesis of ribonucleotides was accompanied by the simultaneous synthesis of arabinonucleotides, 2′-deoxyribonucleotides, and other variations on the canonical nucleotides. In order to understand how relatively homogeneous RNA could have emerged from such complex mixtures, we have examined the properties of arabinonucleotides and 2′-deoxyribonucleotides in nonenzymatic template-directed primer extension reactions. We show that nonenzymatic primer extension with activated arabinonucleotides is much less efficient than with activated ribonucleotides, and furthermore that once an arabinonucleotide is incorporated, continued primer extension is strongly inhibited. As previously shown, 2′-deoxyribonucleotides are also less efficiently incorporated in primer extension reactions, but the difference is more modest. Experiments with mixtures of nucleotides suggest that the coexistence of ribo- and arabino-nucleotides does not impede the copying of RNA templates. Moreover, chimeric oligoribonucleotides containing 2′-deoxy- or arabino-nucleotides are effective templates for RNA synthesis. We propose that the initial genetic polymers were random sequence chimeric oligonucleotides formed by untemplated polymerization, but that template copying chemistry favored RNA synthesis; multiple rounds of replication may have led to pools of oligomers composed mainly of RNA.

---

The RNA world hypothesis proposes that RNA oligonucleotides served dual roles as both genetic and functional polymers during the origin and early evolution of life^1–3^. One of the essential requirements for the emergence of the RNA world is the prebiotic availability of ribonucleotides via abiotic synthesis^4–6^. There has been significant progress in understanding the prebiotic synthesis of the pyrimidine ribonucleotides, led by Sutherland who has demonstrated prebiotically viable syntheses of pyrimidine ribonucleotides (U and C) and their 2-thio variants^7^. Despite recent progress, equivalently direct and high yielding routes to the purine ribonucleotides remain unknown^8,9^. A recent proposal for nucleotide synthesis from the Carrell group is unconstrained with respect to sugar chemistry^10^, suggesting the possibility of a great diversity of nucleotides with differing sugar moieties. Thus, the continued exploration of plausible routes to the canonical ribonucleotides suggests that a diverse set of noncanonical nucleotides may have been present together with ribonucleotides on the early Earth^11,12^. While UV photolysis has been elegantly shown to selectively destroy many undesired side-products in prebiotic pyrimidine nucleotide synthesis^7^, the pool of available nucleotides still remains diverse, with nucleobase variations such as 8-oxo-purine^8^, inosine^8,13^, and the 2-thio-pyrimidines^14^, as well as sugar variants including arabino-^9^, 2′-deoxyribo-^15^, and threo-nucleotides^16,17^.

Intriguingly, some of these variant ribonucleotides have been shown to be compatible with or even beneficial for the copying of RNA templates. For instance, 2-thiouridine greatly improves the rate and fidelity of nonenzymatic primer extension compared with uridine^18^, due to the 2-thio substituent increasing the binding affinity to adenosine and destabilizing wobble pairing with guanosine^19–21^. Inosine, which can arise from the deamination of adenosine, improves the fidelity of nonenzymatic primer extension relative to guanosine^13^. In contrast, the 8-oxo-purine ribonucleotides are extremely poor substrates for nonenzymatic primer extension^13^. Long before recent suggestions of prebiotic routes to the 2′-deoxyribonucleotides^15,22^, these DNA building blocks were known to be less effective than ribonucleotides in nonenzymatic template copying reactions^23^. However, recent work showing that mixed ribo/deoxy oligonucleotide duplexes have much lower melting temperatures than either homogeneous RNA or DNA duplexes^24^ suggests that this heterogeneity might facilitate replication by enhancing strand separation. Given the wide range of effects of noncanonical nucleotides on nonenzymatic template directed copying chemistry, a systematic study of the effects of prebiotically plausible variant nucleotides on copying chemistry is needed in order to understand if and how more homogeneous modern RNA could have arisen from complex prebiotic mixtures of nucleotides.

Among the plausible candidates for primordial nucleosides, arabinonucleosides, the building blocks of arabino-nucleic acid (ANA) are of particular interest (Fig 1a). Powner *et al.* have recently demonstrated the first potentially prebiotic synthesis of all four canonical arabinonucleosides^25^. In this approach, arabino- and ribo-nucleotides derive from the same synthetic intermediate, where external nucleophiles, such as hydroxide or sulfide, furnish the corresponding arabinonucleotides, while an intramolecular 3′ phosphate inverts the stereocenter at the 2′ position to furnish ribonucleotides (Fig 1b). The divergent syntheses of arabino- and ribo-nucleotides from a common intermediate suggests that the first genetic polymer may have been assembled from a heterogeneous mixture of ANA and RNA. Although homogeneous ANA alone cannot form stable duplexes, ANA can form a stable Watson-Crick paired duplex with the corresponding complementary RNA oligonucleotide^26,27^. Recent studies have shown that ANA oligonucleotides can have RNA cleaving catalytic activity when paired across from an RNA oligonucleotide substrate^28^. These results have been taken to suggest that ANA oligonucleotides can serve as both genetic and functional polymers in the presence of RNA oligonucleotides.

**Fig 1:**
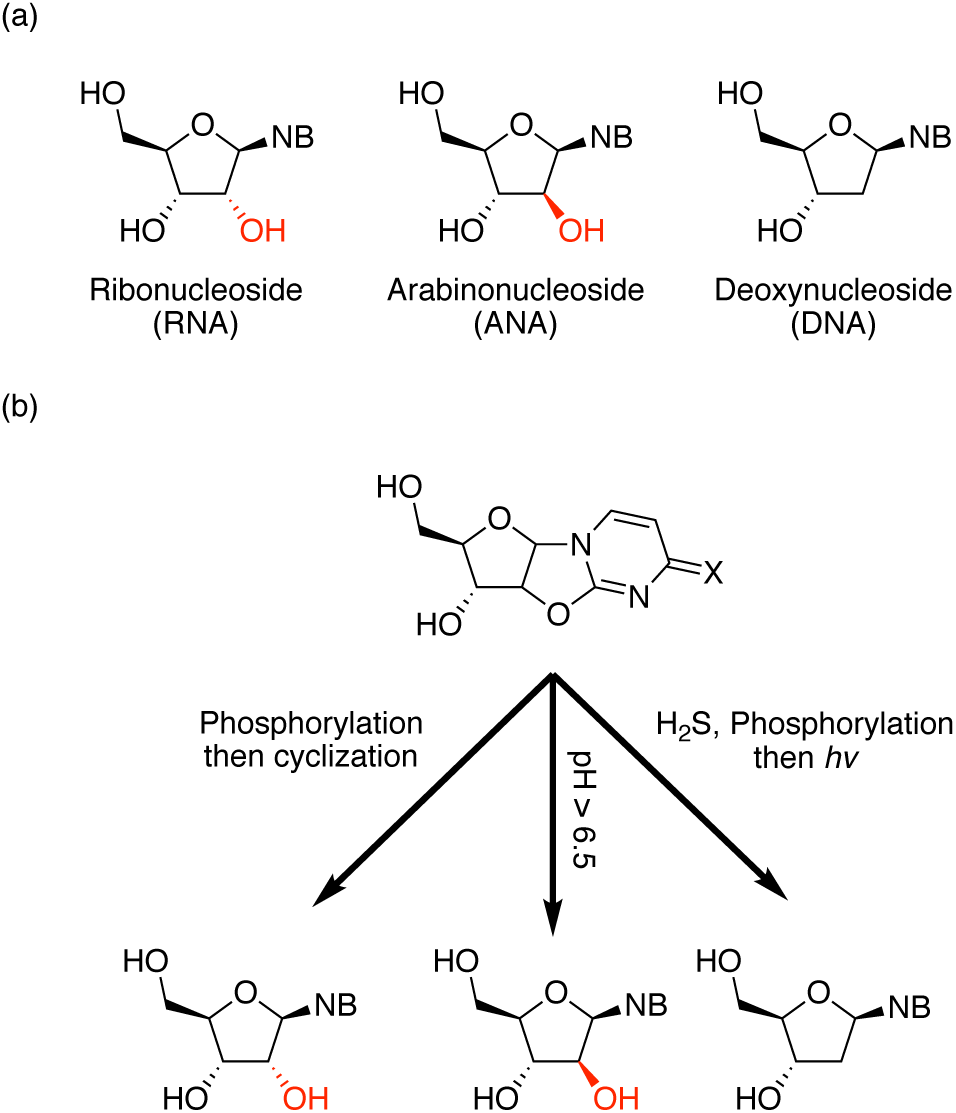
Divergent prebiotic syntheses of nucleosides. (a) Schematic representation of ribo-, arabino-, and 2′-deoxy-nucleosides. NB: nucleobase. (b) Reported divergent syntheses of RNA (left), ANA (middle), and DNA (right) from a common anhydronucleoside intermediate.

Herein, we evaluate the effects of arabinonucleotides on nonenzymatic primer extension, both as activated monomers and when incorporated into primers and templates. We have also revisited the effects of 2′-deoxyribonucleotides in parallel experiments, using more modern and prebiotically plausible activation chemistry than in earlier studies. In addition we have examined the behavior of heterogeneous chimeric, predominantly ribonucleotide oligomers, by systematically introducing single variant residues to the RNA system.

## Results and Discussion

### Primer Extension with Monomers

To examine arabino- and 2′-deoxyribo-nucleotides as substrates in nonenzymatic primer extension, we measured the rate of nonenzymatic primer extension using the 2-aminoimidazole activated monomers, 2AI-araG, 2AI-araA, 2AI-dG, and 2AI-dA across from their respective Watson-Crick pairing partners C and 2^s^U in RNA templates (Fig 2a). We used 2-thio-uridine (2^s^U) instead of uridine in the template because 2^s^U is prebiotically plausible and exhibits stronger pairing with A than U. Previous reports from our lab have demonstrated that activated downstream oligonucleotides can greatly accelerate nonenzymatic primer extensions^29^ by forming highly reactive imidazolium-bridged intermediates^30^ and because of the structural preorganization afforded by an extended helical geometry^31^. Therefore, we prepared 2-aminoimidazole activated RNA trimer helpers and utilized these activated downstream helpers in the evaluation of arabinonucleotides, 2′-deoxyribonucleotides, and ribonucleotides in nonenzymatic primer extension. The rates of primer extension using activated ribonucleotides were the fastest, at 11 hr^−1^ for 2AI-rG and 16 hr^−1^ for 2AI-rA. Primer extension rates with activated 2′-deoxy-nucleotides were slightly slower than with ribonucleotide monomers (8.2 hr^−1^ and 13 hr^−1^ for 2AI-dG and 2AI-dA respectively). In contrast, primer extension with arabinonucleotide monomers was much slower, at 0.88 hr^−1^ for 2AI-araG and 0.52 hr^−1^ for 2AI-araA (Fig 2b).

**Fig 2.**
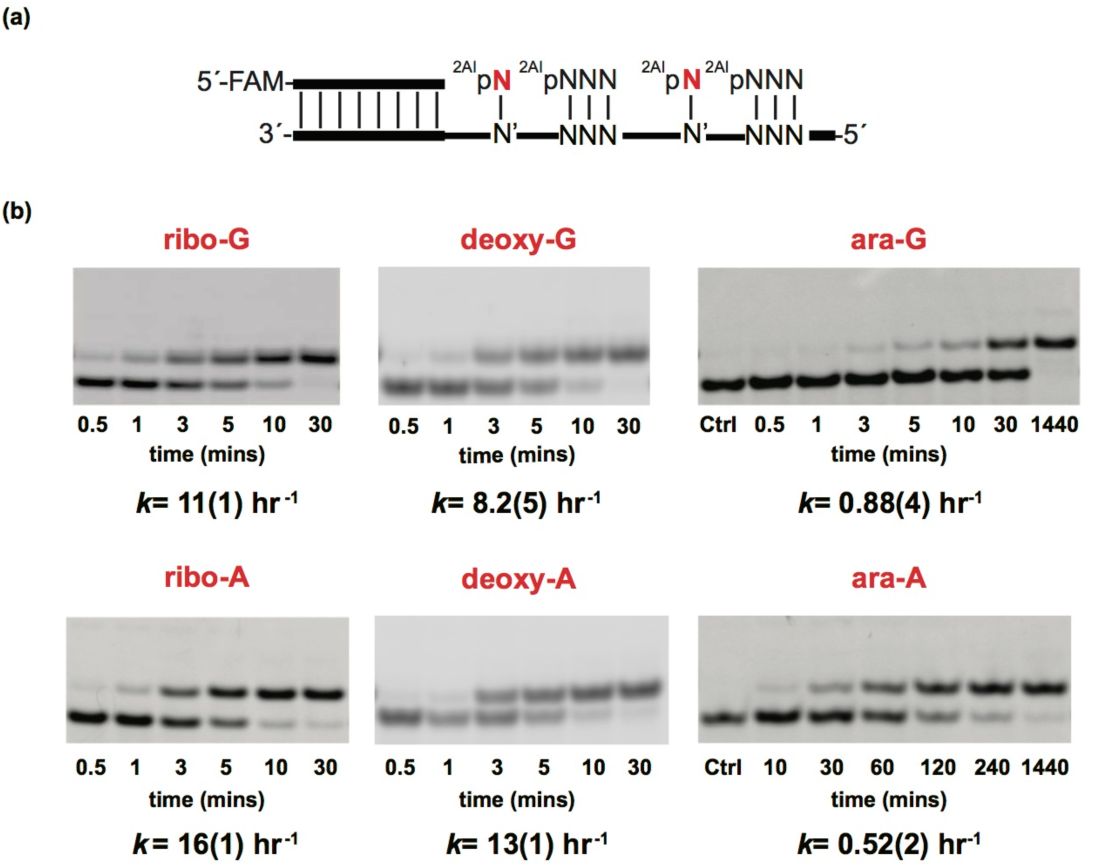
Evaluation of arabino- and 2′-deoxy-nucleotides in nonenzymatic primer extension. (a) Schematic representation of a primer extension reaction using an RNA template. 2AIpN represents 2-aminoimidazole activated monomers and 2AIpNNN represents 2-aminoimidazole activated RNA trimer helpers. (b) Gel electrophoresis images and rates of primer extension for 2-aminoimidazole activated ribonucleotides, 2′-deoxynucleotides, and arabinonucleotides. All reactions were performed at pH 8.0, 200 mM HEPES, 50 mM Mg^2+^, 20 mM 2AIpP_1_, 0.5 mM of activated RNA trimer. See SI appendix for kinetic analysis of primer extension reactions (SI appendix, Fig. S6-S11). Values are reported as the mean ± SD in brackets, with the last digit reported being the last significant figure and the one in which the error arises from triplicate experiments.

In order to distinguish between effects due to the formation, degradation or steady state levels of the imidazolium-bridged intermediate, as opposed to the reactivity of that intermediate once formed, we prepared the 2-aminoimidazolium-bridged dinucleotides from ribo-, 2′-deoxy- and arabino-adenosine. We measured the rates of hydrolysis of the three imidazolium bridged dimers under primer extension conditions (50 mM Mg^2+^, pH 8 in 10% D_2_O; for details see SI Appendix, Fig. S1-S3). Measured rates of hydrolysis revealed no significant difference in the stability of activated dimer species with *k*= 0.31 hr^−1^, 0.33 hr^−1^ and 0.30 hr^−1^ for rA dimer, araA dimer and dA dimer respectively, showing that the stereochemical information at the 2′ position does not impact the decomposition of the dimer intermediate under relevant conditions. In our evaluation of the rate of dimer intermediate formation, ribonucleotides, arabinonucleotides and deoxynucleotides show no meaningful difference with *k*= 0.0032 hr^−1^ mM^−1^, 0.0031 hr^−1^ mM^−1^ and 0.0028 mM^−1^ for rA, araA and dA respectively (see SI Appendix, Fig. S4-S5).

We then evaluated the Michaelis-Menten parameters for primer extension for all three imidazolium-bridged dinucleotides. For these experiments, we prepared RNA templates containing two consecutive s^2^U residues by solid phase synthesis; we also used downstream oligonucleotides to further promote the nonenzymatic copying (Fig 3a). The *K*_m_ values of the imidazolium-bridged dimer intermediates are very low, at 0.11 mM, 0.065 mM, and 0.64 mM for rA, dA, and araA respectively (Fig 3c). While these *K*_m_ values span an order of magnitude, all values, including *K*_m_ for araA, are low, in line with the ability to form ANA:RNA duplexes^24^. Interestingly, *V*_max_ for the three dimer intermediates differ by approximately 70 fold, at 22 mM hr^−1^, 1.4 mM hr^−1^ and 0.32 mM hr^−1^ for rA, dA, and araA respectively (Fig 3c). The large differences in *V*_max_ are surprising, since the 2′-substituent is remote from the site of the reaction between the primer 3′-hydroxyl and the phosphate of the adjacent incoming nucleotide. Furthermore, simple nucleophiles such as hydroxide or water attack the phosphate of all three imidazolium-bridged dimers with similar rates, indicating that the 2′-substituent influences the rate of phosphate substitution only with stereochemically rich nucleophiles, such as the ribose at the end of the primer. Arabino-nucleotides have been shown to adopt a C2′ endo pucker with intramolecular hydrogen bonding between the 2′-hydroxyl and 5′-oxygen^32^, while 2′-deoxynucleotides show a more modest preference for a C2′ endo pucker and ribonucleotides adopt a C3′ endo pucker. This difference in sugar conformation could affect the position of the 5′-phosphate of the nucleotide adjacent to the 3′-end of the primer, potentially explaining the observed rate differences.

**Fig 3.**
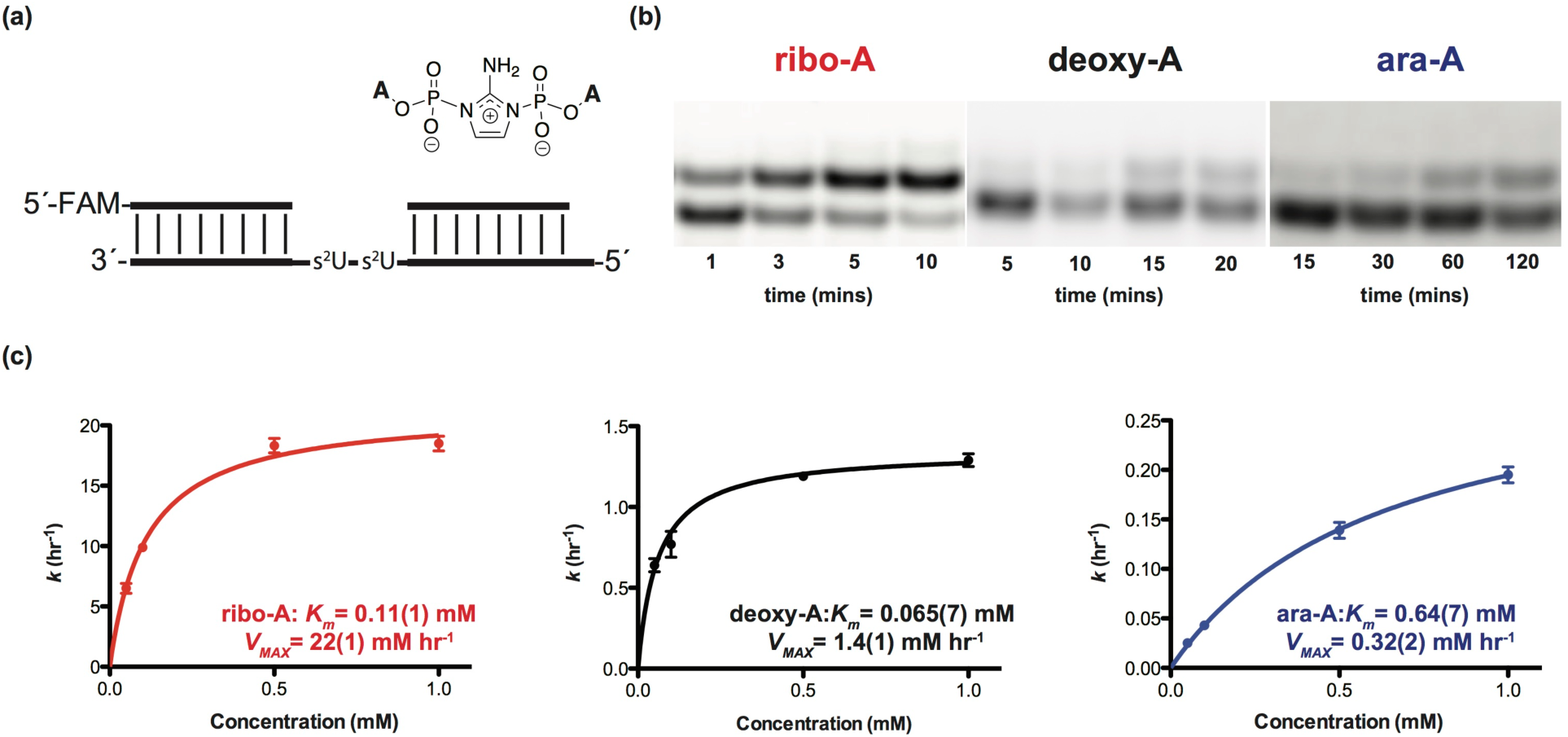
Michaelis-Menten study of nonenzymatic copying using rA, dA, and araA imidazolium bridged dimers. All reactions were performed at pH 8.0, 200 mM HEPES, 50 mM Mg^2+^, and various concentrations of homo-dimers ranging from 0.05 to 1 mM. (a) Schematic representation of a primer extension reaction with imidazolium bridged dimers. (b) Gel electrophoresis images of primer extension for imidazolium bridged dimers at 1 mM concentration. (c) Plot of *k* (hr^−1^) against the concentration of rA, dA, and araA imidazolium bridged dimers. See SI appendix for kinetic analysis of primer extension reactions (SI appendix, Fig. S12-S26). Values are reported as the mean ± SD in brackets, with the last digit reported being the last significant figure and the one in which the error arises from triplicate experiments.

### Effect of arabino- and 2′-deoxy-nucleotides at the 3′-end of the primer

Previous studies indicate that the nature of the 3′-end of the primer can strongly influence the rate of subsequent nonenzymatic primer extension^33^; up to a 300 fold decrease in the rate of primer extension has been observed for a mismatched base pair vs. a Watson-Crick base pair at the 3′-end of the primer^13^. We therefore evaluated the effect on nonenzymatic primer extension of replacing the ribonucleotide at the 3′-end of the primer with either a 2′-deoxynucleotide or an arabinonucleotide. We prepared the three primers by solid phase synthesis, and then measured the rate of primer extension in the presence of 2-aminoimidazole activated guanosine monomers using templates containing the Watson-Crick paired primer binding region followed by a C_3_ templating region (Fig 4a). Strikingly, primers containing a single arabinonucleotide at the 3′-end showed no measurable extension even after 4 hours. In contrast, primers with ribo- or 2′-deoxy-nucleotides at the 3′-end were rapidly extended under the same reaction conditions (*k*_obs_= 4.6 hr^−1^ and 2.8 hr^−1^ for primers containing a 3′ terminal rG and rA *vs.* 2.1 hr^−1^ and 1.3 hr^−1^ for dG and dA, respectively), showing that the change in the configuration of the 2′-hydroxyl group on the pentose at the 3′ end of primer strongly affects nonenzymatic primer extension.

**Fig 4.**
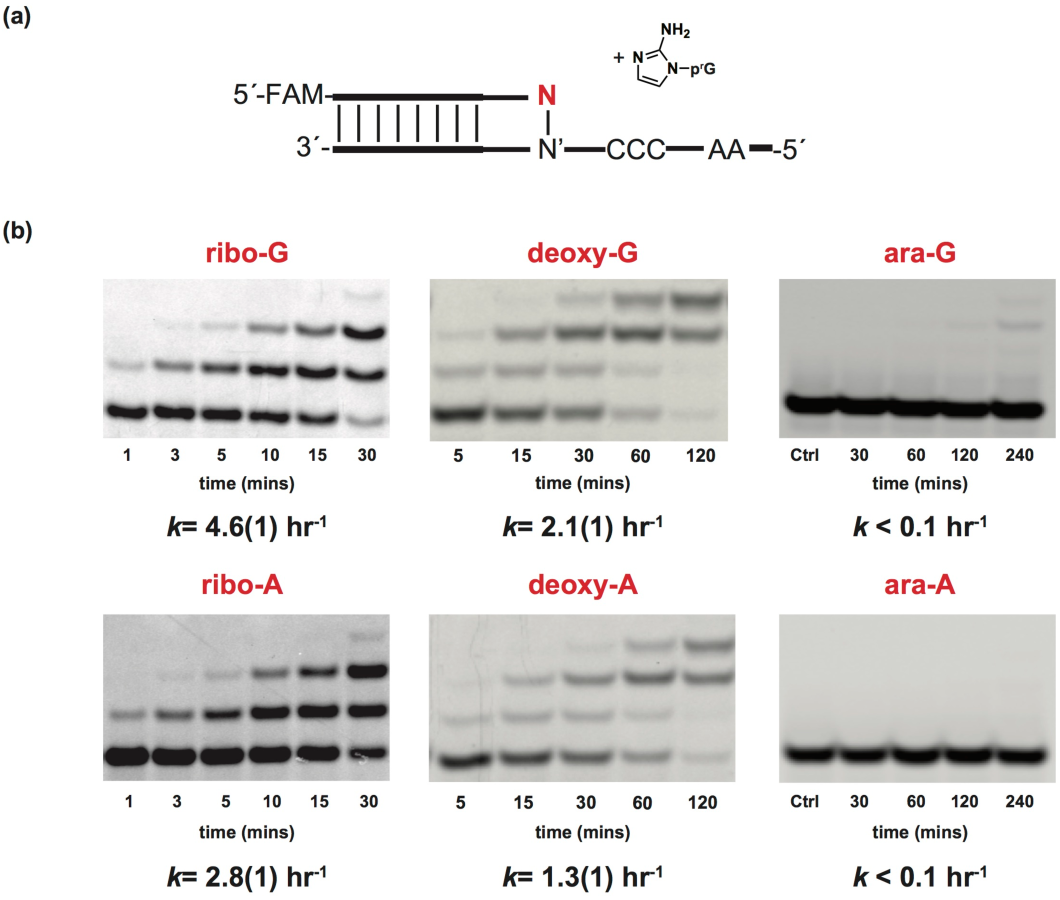
Evaluation of nonenzymatic primer extension with primers containing alternative nucleotides at the 3′ end. (a) Schematic representation of nonenzymatic primer extension of 2AIrG with a primer that contains either a ribo-, a 2′-deoxy-, or an arabino-nucleotide (P_0_) at the 3′ end. (b) Gel electrophoresis images and rates of primer extension for primers containing ribo-, 2′-deoxy-, and arabino-nucleotides at the 3′ end. See SI appendix for kinetic analysis of primer extension reactions (SI appendix, Fig. S27-S32). Values are reported as the mean ± SD in brackets, with the last digit reported being the last significant figure and the one in which the error arises from triplicate experiments.

Given the dramatic stalling effect of an arabinonucleotide at the 3′ end of the primer, we sought to understand the reason for this effect at the atomic level. We therefore crystallized a chimeric oligonucleotide with a 3′-arabinonucleotide, complexed with RNA-monomers. The rGMP monomer was co-crystallized with a self-complementary 14mer RNA 5′-**^m^C^m^C^m^CG**ACUUAAGUCaG-3′, which contains 4 locked nucleotides at the 5′ end (in bold) to rigidify the template and speed up crystallization (Fig 5a). The residue at the 3′ end is arabino-G (_a_G), to mimic the nonenzymatic primer extension shown in Fig 4. At each end, two GMP nucleotides bind to the templating locked 5-methyl Cs, forming a 16mer RNA-monomer helical structure. The assembled complex crystallizes in the rhombohedral R32 space group, and the structure was determined to 2.0-Å resolution. In one asymmetric unit, one strand of RNA is complexed with 2 GMP monomers. Except for the crystallographic symmetry, this arabino-modified RNA complex structure resembles our previously determined RNA-GMP complex structures^34,35^, including the A-form double helices, C3′-endo conformation of the RNA sugars and slip-stacking of the neighboring duplexes (Fig 5b). The crystal structure reveals two key features of the terminal arabino-G residue: it is in the C2′-endo conformation, and forms a Watson-Crick base pair with the templating ^**m**^**C**. The sugar of the terminal arabino-G is well-ordered, and its C2′-endo conformation contrasts with reported structures of an RNA/ANA heteroduplex, in which the arabinonucleotides are in the O4′-endo (Eastern) conformation ^24,26,36^ (Fig 5c). The nucleobase of the arabino-G residue is Watson-Crick base paired with the templating ^**m**^**C**, with hydrogen bonds ranging from 2.8 to 3.1 Å.

**Fig 5.**
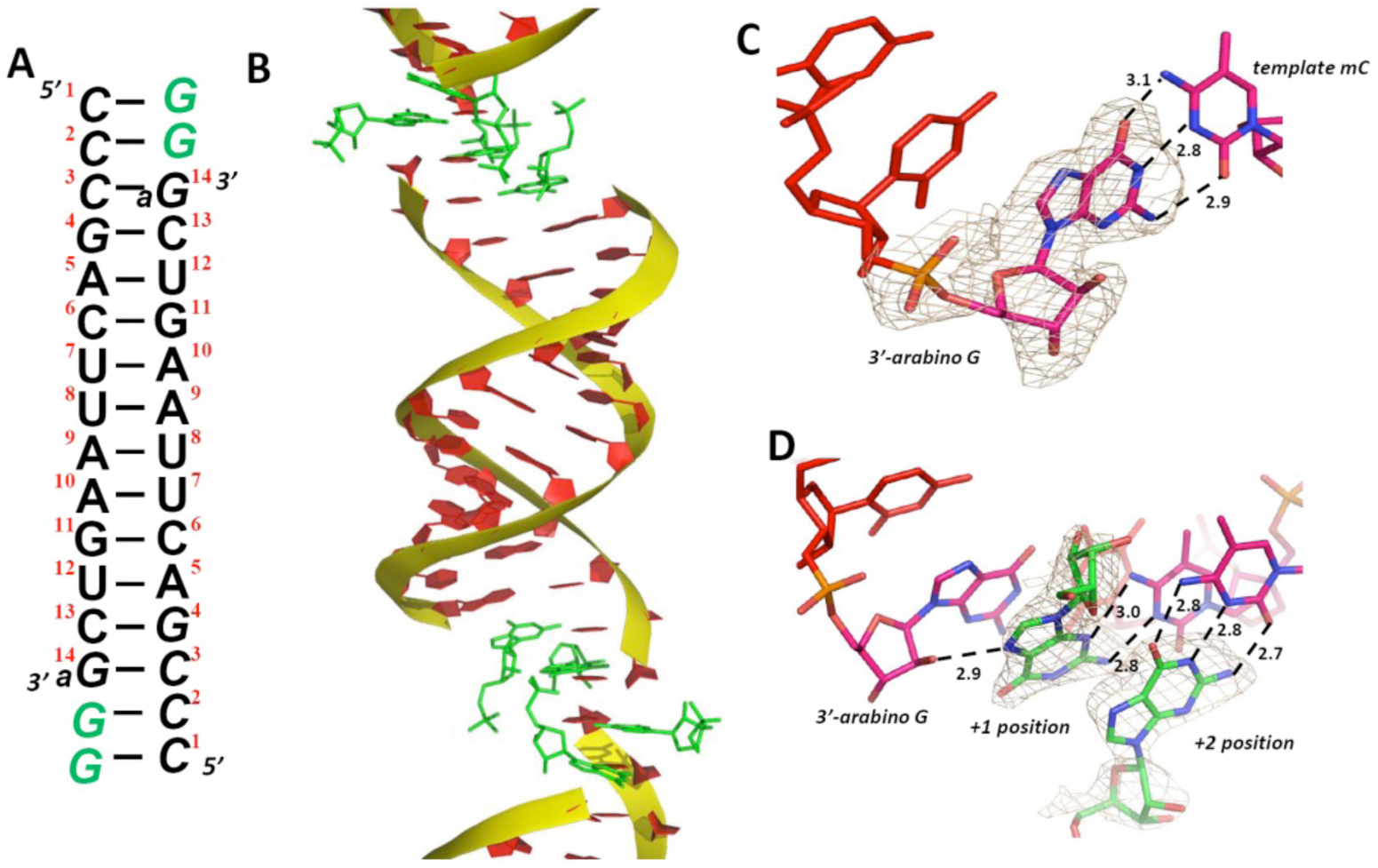
Crystal structure of RNA primer/template complex with an arabinonucleotide at the 3′-end of the primer (PDB 6OWL). Red: primer; green: GMP monomers. The wheat mesh indicates the 2*F*_*o*_ − *F*_*c*_ omit map contoured at 1.5σ. (A) Schematic of the primer/template complex. Italic letters: locked nucleotides; _a_G at the 3′ end: arabino-guanosine. (B) Overall structure of the 3′-arabino RNA-GMP complex, stacking with neighboring complexes. (C) The local structure of the arabino-G-terminated primer. The corresponding omit map indicates that the primer arabinose is in the C2′-endo conformation, and the guanine is Watson-Crick base paired with the template. (D) The GMP monomers are bound through both canonical and non-canonical G:C base pairs. The primer 2′-OH and N7 of the +1 GMP monomer are within hydrogen bonding distance.

In the evaluation of various nucleotides in nonenzymatic primer extension, several prior reports have shown a strong correlation between primer extension rates and the conformation of the 3′-terminal nucleotide. Primers composed of or ending in nucleotides that prefer a C3′-endo conformation (A-form) engage in faster primer extension than primers composed of or ending in nucleotides that do not adopt a C3′-endo conformation (e.g. DNA)^23,37,38^. Additionally, pseudoaxial/axial nucleophiles generally exhibit lower reactivities than pseudoequatorial/equatorial nucleophiles^39^. This is due to steric encumbrance of axial/pseudoaxial substituents, especially by 1,3-diaxial interactions, which are exacerbated by a more crowded steric environment in the product and in the transition state. One plausible explanation for the significant stalling effect we observe in nonenzymatic polymerization with an arabino-nucleotide terminated primer is that the preferred C2′-endo conformation of the arabino-nucleotide places the 3′-hydroxyl too far away from the phosphate of the incoming monomer.

The base-pairing of the unactivated GMP monomer bound downstream of the primer is significantly different from that observed in our previously determined RNA-GMP structures. A hydrogen bonding interaction between the N7 atom of the +1 GMP and the 2′-OH group of the arabino-nucleotide at the end of the primer is observed (2.9 Å, Fig 5d). This flips the nucleobase of the +1 GMP into a non-canonical G:C base pair in which N3 of the +1 GMP forms a hydrogen bond with the exocyclic amine of the templating ^**m**^**C**, while the exocyclic amine of the guanine hydrogen bonds with N3 of the ^**m**^**C**. This noncanonical base pairing positions the 5′-phosphate of the monomer distant from the 3′-OH of the primer. The phosphate of the +1 GMP is highly disordered, and the primer 3′-OH of the arabinose sugar is pointing to the minor groove instead of the incoming monomer. In previously determined RNA-GMP structures, the first bound GMP monomer was Watson-Crick paired with the template, with its binding affinity and the distance between the primer 3′-OH and the incoming phosphorus improved by the downstream bound monomer at the +2 position. The structural changes observed in the case of the arabino-terminated primer would be detrimental to nonenzymatic primer extension, in line with our experimental data; however the crystal structure may not reflect the conformation of the template bound imidazolium-bridged dinucleotide.

### Competition Experiment between Activated rA and araA

The common intermediate in a recently proposed prebiotic synthesis of RNA, DNA, and ANA raises the possibility that these nucleotides may have coexisted on early earth^25^. Given the strong stalling effect when the 3′ end of the primer is an arabinonucleotide, the incorporation of an arabinonucleotide into a growing primer will essentially act as a chain terminator, potentially preventing the synthesis of oligonucleotides long enough to have catalytic function. Fortunately, arabino-nucleotide incorporation rates were 15-30 fold slower than ribonucleotide incorporation rates in the case of trimer assisted nonenzymatic copying, suggesting that ribonucleotides may outcompete arabinonucleotides well enough to minimize this concern. To test this idea, we performed a nonenzymatic copying reaction in a one pot competition experiment with a 1:1 ratio of activated araA and rA monomers, utilizing the reaction conditions in Figure 2. The initial primer, P1, was exposed to activated rA and araA monomers with activated trimer helper (2AI-rGAC) for one hour, to allow for incorporation of either rA (generating primer P2) or araA (generating primer P3). A subsequent nonenzymatic copying step was then performed by adding the activated RNA monomer 2AI-rG and the trimer helper 2AI-rACA. Primers that initially incorporated araA to generate P3 cannot be extended in the second step (Lane 7, Fig 6b) while primers that initially incorporated rA to generate primer P2 can be further extended (Lane 5, Fig 6b). After 4 hours, the reaction was quenched and PAGE gel analysis was performed to analyze the product ratios (Fig 6b). Only a small fraction (approximately 7%) fails to extend after the initial addition (Lane 3, Fig 6b). Generally, iterative nonenzymatic copying of oligonucleotides is less than 100% complete^29,40^, presumably due to the presence of multiple trimer helpers, which have high binding affinity to the template and are thus inhibitory to subsequent copying. Our results therefore represent a lower bound on the true ratio of rA vs. araA incorporation, and indicate that activated rA monomers substantially outcompete activated araA in the nonenzymatic copying of oligonucleotides. Thus, the coexistence of arabino- and ribo-nucleotides should not significantly interfere with oligonucleotide copying.

**Fig 6.**
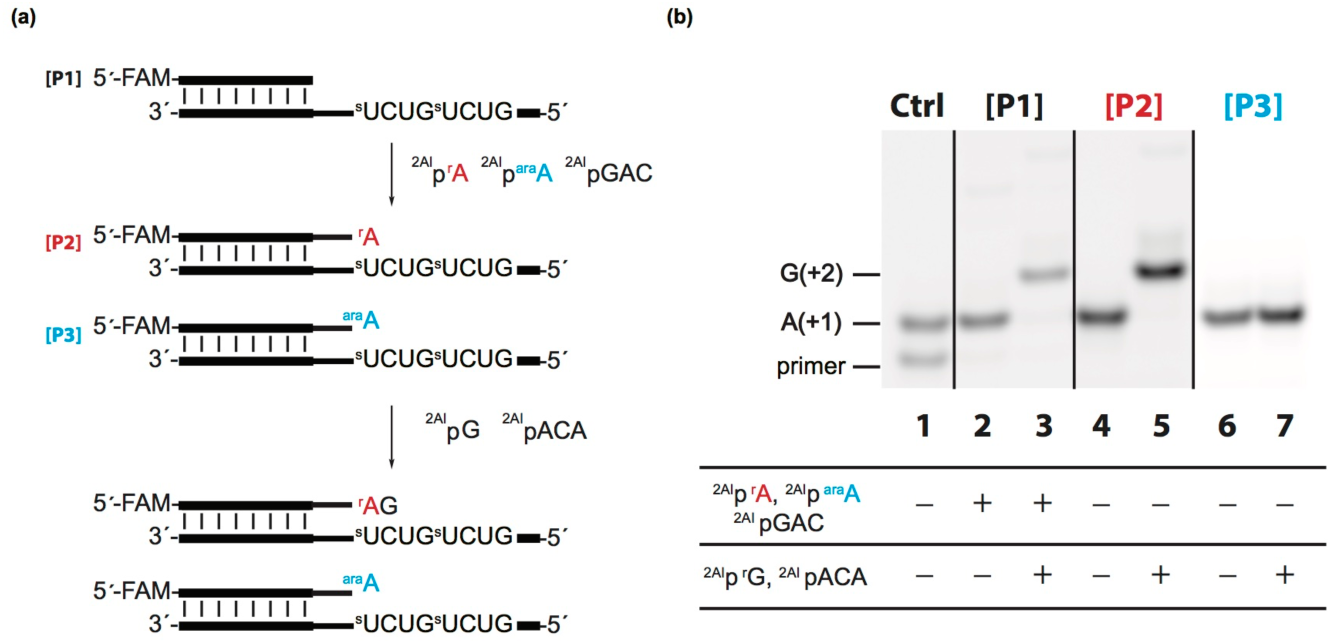
Competition experiment between 2AI-araA and 2AI-rA. (a) Schematic representation of nonenzymatic primer extension with both ribonucleotide and arabinonucleotide monomers followed by a nonenzymatic copying step, in which we expect the ribo-terminated primer [P2] to extend and the arabino-terminated primer [P3] not to extend (shown in lanes 5 and 7 in b). (b) Gel electrophoresis images of the competition experiment. Primers used for the experiment are shown on the top ([P1] for lanes 2/3, [P2] for lanes 4/5, and [P3] for lanes 6/7) and the activated nucleotide monomers and trimer helpers added are shown below. Synthetic primers [P2] in lane 4 and [P3] in lane 6 were used as controls. Ctrl lane 1 includes synthetic primers [P1] and [P2].

### Primer Extension Across an Arabinonucleotide in the Template Strand

Although the above results suggest that arabinonucleotide-containing oligonucleotides would only rarely be generated by nonenzymatic primer extension, untemplated polymerization of activated monomers could lead to the synthesis of oligonucleotide templates containing arabinonucleotides in internal positions ^41 42^. We have evaluated the effects of internal arabinonucleotides in template strands on the rate of primer extension with standard activated ribonucleotides. We prepared RNA templates containing isolated arabino- or 2′-deoxy-nucleotides by solid phase synthesis, and then performed nonenzymatic primer extensions with activated RNA monomers and trimer helpers, and compared the observed rates on all-RNA templates and RNA templates containing single arabino- or 2′-deoxynucleotide residues (Fig 7a). Primer extension with 2-aminoimidazole activated cytidine exhibited a minimal rate difference when copying across rG, dG and araG in the template strand, with *k*_obs_= 10, 8.0, and 6.8 hr^−1^ (Fig 7b). Primer extension with activated 2-thiouridine also exhibited minor rate differences when copying across rA, dA and araA in the template strand, with *k*_obs_= 12, 11, and 2.1 hr^−1^, respectively. In all cases, we observe complete conversion of the primer to extended products in the presence of the corresponding complementary Watson-Crick RNA monomers.

**Fig 7.**
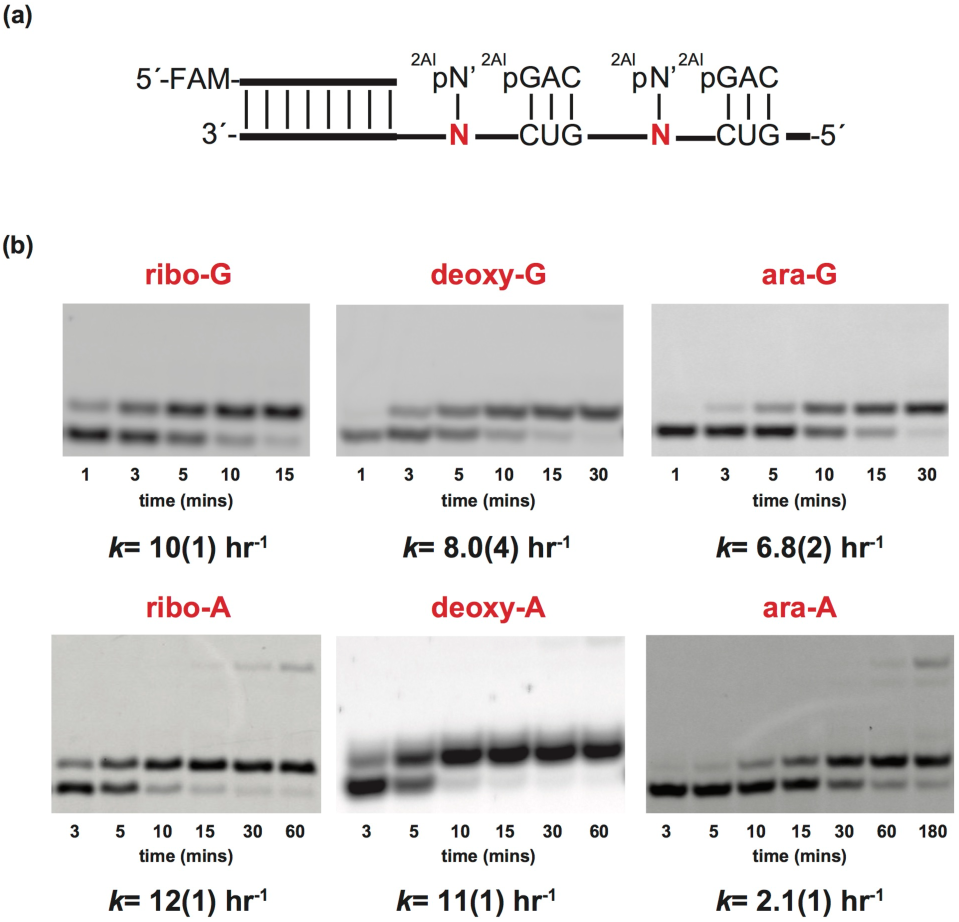
Evaluation of primer extension for 2-aminoimidazole activated pyrimidine nucleotides across from templating ribo-, 2′-deoxy-, and arabino-nucleotides. (a) Schematic representation of a primer extension reaction with template containing a ribo-, 2′-deoxy-, or arabino-nucleotide. 2AIpN represents the corresponding 2-aminoimidazole activated ribonucleotide monomers and 2AIpNNN represents 2-aminoimidazole activated RNA trimer helper. (b) Gel electrophoresis images and rates of primer extension for 2-aminoimidazole activated pyrimidine ribose across from a ribo-, 2′-deoxy- and arabino-nucleotide. See SI appendix for kinetic analysis of primer extension reactions (SI appendix, Fig. S33-S38). Values are reported as the mean ± SD in brackets, with the last digit reported being the last significant figure and the one in which the error arises from triplicate experiments.

## Conclusions

The discovery of potentially prebiotic synthetic routes to the canonical ribonucleotides has given rise to an improved understanding of the most likely alternative nucleotide byproducts of these pathways. The likely presence of these byproducts leads to a significant problem with regard to the emergence of the RNA World, since the initially synthesized oligonucleotides would be expected to be quite heterogeneous in composition. How could such a heterogeneous mixture of oligonucleotides give rise to the relatively homogeneous RNAs that are thought to be required for the evolution of functional RNAs such as ribozymes?

In addressing the problem of the emergence of RNA, we have, in this paper, focused on the consequences of co-existing activated arabino- and 2′-deoxy-nucleotides for nonenzymatic template directed primer extension. We find that both primer extension with activated arabino-nucleotides, and the extension of an arabinonucleotide-terminated primer with ribonucleotides, are very slow. Our one pot competition experiment between ribo- and arabino-nucleotides directly demonstrates that highly RNA enriched oligonucleotides could arise by template copying with a heterogeneous mixture of nucleotides. Our kinetic and structural studies are consistent with this rate difference being due to the 2′-endo sugar conformation of arabinonucleotides. In parallel experiments, we find, as expected^23^, that primer extension with 2′-deoxynucleotides, and extension of a 2′-deoxynucleotide terminated primer, are slower than with ribonucleotides, but the differences are much smaller than in the case of arabino-nucleotides. In contrast, the copying of templates containing either an arabino- or a 2′-deoxynucleotide is only moderately slower than the copying of an all RNA template. Our results suggest that an initial set of highly heterogeneous oligonucleotides, generated by non-templated polymerization, could act as templates for the synthesis of complementary strands that would be enriched in RNA. Multiple cycles of template copying by primer extension could therefore provide a pathway for a transition from heterogeneity to homogeneity. We have seen similar effects in previous work with 8-oxo-purine ribonucleotides, where primer extension with these activated nucleotides is extremely slow, but they can be copied when present in the template strand, albeit at lower rates^13^.

The Krishnamurthy group has recently proposed a similar pathway leading from an initially heterogeneous set of templates to an increasingly homogeneous set of replication products^43^. In their case the physical mechanism leading to homogeneity is different, and is based on the surprising observation that homogeneous all-RNA or all-DNA oligonucleotides bind more strongly than do otherwise identical but heterogeneous oligonucleotides to complementary but heterogeneous long templates. As a result of the stronger binding, the template-directed ligation of all RNA or all DNA oligonucleotides is favored over the ligation of mixed oligonucleotides, such that over repeated cycles of ligation-mediated replication, relatively homogeneous replication products should emerge. Because these experiments relied on non-prebiotic ligation chemistry and the nucleotide variation was restricted to internal positions, effects on ligation rates could not be evaluated. However variations in chemical reactivity might also be expected to contribute to the emergence of homogeneous products. We have previously suggested that both primer extension with activated monomers and ligation mediated template copying may have acted together to mediate primitive replication cycles^44^, in which case both ligation and primer extension effects may have contributed to the emergence of homogeneous RNA oligonucleotides as a result of cycles of replication. We note that both primer extension with monomers and oligonucleotide ligation are driven by the energy of nucleotide (or oligonucleotide) activation. The chemical energy of activated substrates drives template copying and ultimately replication in a far from equilibrium reaction network, which in turn powers the emergence of homogeneity from heterogeneity. Thus, local order in the form of homogeneous oligonucleotides emerges at the expense of chemical energy that is dissipated through an overall increase in disorder.

Our observations show that arabino-terminated products of primer extension cannot be further extended at significant rates. In the past, such dead-end products of incomplete replication would have been considered to be highly detrimental to the overall process of genomic replication in protocells. However, recent results suggest that such oligonucleotides could in fact have multiple positive or even essential roles in primordial genomic replication. For example, when suitably activated, short oligonucleotides act as catalysts of template copying by forming imidazolium-bridged intermediates with activated monomers^29,30^. Slightly longer oligonucleotides may also act as catalysts of strand displacement synthesis through a toehold/branch migration mechanism that assists in opening up a template and allowing primer extension to proceed^45^. Similar oligonucleotides could also act as splint templates for nonenzymatic ligation; since they could not be elongated by primer extension they could be much easier to dissociate from the longer ligated product, which could facilitate the construction of functional oligonucleotides^46^. Short oligonucleotides may also contribute to homeostatic regulation of ribozyme activity in protocells^47^.

Multiple distinct processes, in addition to primer extension and ligation, are likely to have contributed to the transition from heterogeneous primordial nucleic acids to the relatively homogeneous RNA genomes of the first cells. For example, photochemical reactions preferentially degrade the noncanonical nucleobases (and the corresponding nucleosides and nucleotides) and also preferentially degrade the alpha-anomeric byproducts of nucleotide synthesis^7^. Steric constraints and variations in chemical reactivity may have influenced the composition of the first oligonucleotides formed through non-templated polymerization; for example, nucleotides with acyclic sugars rapidly cyclize to unreactive products following phosphate activation. The synthesis of standard 3′-5′ phosphodiester bonds when copying templates with 2′-5′ linkages^48^ or 3′-5′ pyrophosphate linkages^49^ may also have contributed to the gradual elimination of the variability in nucleic acid structure that is the inevitable consequence of nonenzymatic polymerization. Taken together these and other mechanisms may explain the transition from a heterogeneous mixture of prebiotically synthesized nucleotides and oligonucleotides to a relatively homogeneous set of RNAs. Considerable additional experimental work must be done to extend this model, as only a fraction of the likely prebiotic variability in nucleotide and nucleic acid structure has been explored to date.

## Acknowledgements

J.W.S. is an Investigator of the Howard Hughes Medical Institute. This work was supported in part by a grant (290363) from the Simons Foundation to J.W.S. and by a grant from the NSF (CHE-1607034) to J.W.S. D.K.O. is a recipient of a Postdoctoral Research Scholarship (B3) from the Fonds de recherche du Québec – Nature et technologies (FRQNT), Quebec, Canada, and a Postdoctoral Fellowship from Canadian Institutes of Health Research (CIHR) from Canada. The authors would like to thank Andrew J. Bendelsmith, Li Li, and Victor S. Lelyveld for their helpful discussions. The authors also thank the entire Szostak group for feedback.

## TOC Graphics

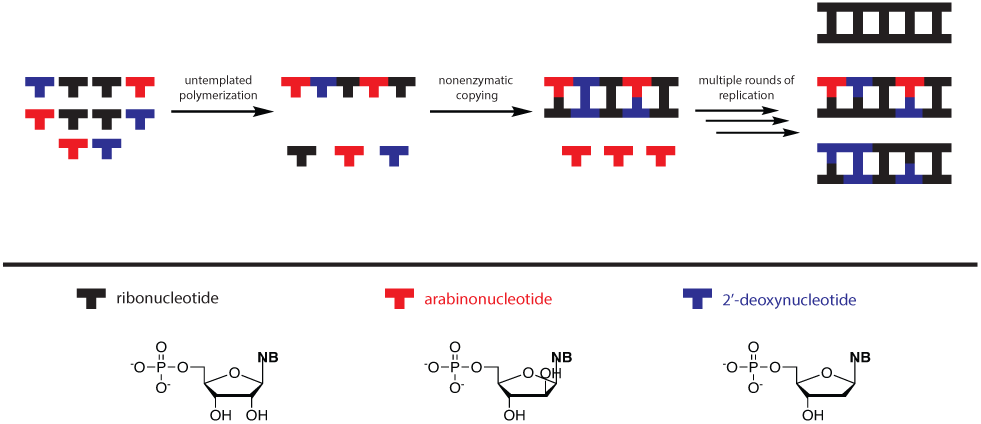

