## Supplemental Info for "A model for the emergence of RNA from a prebiotically plausible mixture of ribonucleotides, arabinonucleotides and 2’-deoxynucleotides"

**This PDF file includes:**

Materials and Methods
Figs. S1 to S40
References for SI reference citations

**Table for Contents**

**Materials and Methods**


**Oligonucleotide synthesis……………………………………………………………3**

**Synthesis of activated nucleotides…………………………………………………3**

**Hydrolysis of activated araA and rA imidazolium bridged dimers (Figures S1-S4)……………………………………………………………………………5**

**Formation of activated araA and rA imidazolium bridged dimers (Figures S5-S7)……………………………………………………………………………9**


**Supporting Figures**


**SI Figures for "Evaluation of arabino- and 2'-deoxy- nucleotides in nonenzymatic primer extension (Fig 2)" (Figures S8 to S13)……………………………………………………………………12**

**SI Figures for "Michaelis-Menten study of imidazolium intermediates of rA, dA, and araA imidazolium bridged dimers (Fig 3)" (Figures S14 to S28)……………………………………………………………………18**

**SI Figures for "Evaluation of nonenzymatic primer extension with primers containing various pentose nucleotides at the 3' end (Fig 4)" (Figures S29 to S34)……………………………………………………………………33**

**SI Figures for "Evaluation of primer extension for 2-aminoimidzole activated pyrimidine nucleotides across various pentose nucleotides (Fig 7)" (Figure S35 to S40)……………………………………………………………………39**

**References……………………………………………………………………………44**

### Materials and Methods

#### 1.1 Oligonucleotide Synthesis

All oligonucleotides used in this study were purchased from Integrated DNA Technologies (Coralville, IA) or prepared by solid-phase synthesis using an Expedite 8909 DNA/RNA synthesizer. Synthesizer reagents and phosphoramidites were purchased from Glen Research (Sterling, VA) and Chemgenes (Wilmington, MA). In-house prepared oligonucleotides were deprotected using standard methods and subsequently purified by polyacrylamide gel electrophoresis. The oligonucleotides were analyzed by high resolution mass spectrometry (HRMS) on an Agilent 6520 QTOF LC-MS.

#### 1.2 Synthesis of activated nucleotides

2AIparaA, 2AIparaG, 2AIpdA, 2AIpdG, 2AIprA, and 2AIprG were prepared according to a previously reported procedure[1]. GAC and AGG 5′-phosphoro-2-aminoimidazolide were prepared as described previously[1]. 2AIprA-imidazolium bridged dimer, 2AIpdA-imidazolium bridged dimer, and 2AIparaA-imidazolium bridged dimer were prepared as described previously[2]. All nucleotides were purified using reverse phase chromatography using a Teledyne Isco Combiflash on a RediSepRf C18Aq column with 2 mM triethylammonium bicarbonate (pH 7.5) and acetonitrile as eluents.

((2R,3R,4R,5R)-5-(6-amino-9H-purin-9-yl)-3,4-dihydroxytetrahydrofuran-2-yl)methyl hydrogen (2-amino-1H-imidazol-1-yl)phosphonate (2AIparaA): $^{1}$H NMR (400 MHz, Deuterium Oxide) δ 8.17 (s, 1H), 8.05 (s, 1H), 6.69 (t, J = 1.7 Hz, 1H), 6.46 (t, J = 2.1 Hz, 1H), 6.37 (d, J = 6.0 Hz, 1H), 4.59 (d, J = 6.2 Hz, 1H), 4.42 (t, J = 6.6 Hz, 1H), 4.22 – 4.14 (m, 2H), 4.14 – 4.08 (m, 1H). $^{31}$P NMR (162 MHz, Deuterium Oxide) δ -10.59. HRMS: calc for [$C_{13}H_{16}N_8O_6P^-$] 411.0936, found 411.0933.

((2R,3R,4R,5R)-5-(2-amino-6-oxo-5,6-dihydro-9H-purin-9-yl)-3,4-dihydroxytetrahydrofuran-2-yl)methyl hydrogen (2-amino-1H-imidazol-1-yl)phosphonate (2AIparaG): $^{1}$H NMR (400 MHz, Deuterium Oxide) 7.88 (s, 1H), 6.68 (t, *J* = 1.7 Hz, 1H), 6.45 (t, *J* = 2.2 Hz, 1H), 6.17 (d, *J* = 6.1 Hz, 1H), 4.54 (t, *J* = 6.3 Hz, 1H), 4.45 (t, *J* = 6.6 Hz, 1H), 4.26 – 4.14 (m, 2H), 4.14 – 4.00 (m, 1H). $^{31}$P NMR (162 MHz, Deuterium Oxide) δ -10.71. HRMS: calc for [$C_{13}H_{17}N_8O_7P\text{-}H^+$] 429.1031, found 429.1046.

((2S,4R,5R)-5-(6-amino-9H-purin-9-yl)-4-hydroxytetrahydrofuran-2-yl)methyl hydrogen (2-amino-1H-imidazol-1-yl)phosphonate (2AIpdA): $^{1}$H NMR (400 MHz, Deuterium Oxide) δ 8.28 (s, 1H), 8.23 (s, 1H), 6.54 (dd, J = 2.3, 1.7 Hz, 1H), 6.48 – 6.38 (m, 2H), 4.25 (d, J = 5.4 Hz, 1H), 4.11 (dd, J = 5.5, 2.8 Hz, 2H), 2.91 (dt, J = 13.7, 6.3 Hz, 1H), 2.62 (ddd, J = 14.2, 7.4, 4.6 Hz, 1H). $^{31}$P NMR (162 MHz, Deuterium Oxide) δ -10.02. HRMS: calc for [$C_{13}H_{16}N_8O_5P^-$] 395.0987, found 395.0982.

((2S,4R,5R)-5-(2-amino-6-oxo-5,6-dihydro-9H-purin-9-yl)-4-hydroxytetrahydrofuran-2-yl)methyl hydrogen (2-amino-1H-imidazol-1-yl)phosphonate (2AIpdG): $^{1}$H NMR (400 MHz, Deuterium Oxide) δ 7.93 (s, 1H), 6.61 (t, J = 1.8 Hz, 1H), 6.46 (t, J = 2.0 Hz, 1H), 6.27 (t, J = 6.7 Hz, 1H), 4.68 – 4.62 (m, 1H),4.20 (d, J = 4.0 Hz, 1H), 4.07 (t, J = 4.8 Hz, 2H), 2.86 (dt, J = 13.7, 6.5 Hz, 1H), 2.51 (ddd, J = 14.2, 6.8, 4.3 Hz, 1H). $^{31}$P NMR (162 MHz, Deuterium Oxide) δ -10.32. HRMS: calc for [$C_{13}H_{17}N_8O_6P\text{-}H^+$] 413.1081, found 413.1085.

Bis(((2R,3S,4R,5R)-5-(6-amino-9H-purin-9-yl)-3,4-dihydroxytetrahydrofuran-2-yl)methyl) (2-amino-1H-imidazole-3-ium-1,3-diyl)bis(phosphonate) (2AIprAimidazolium bridged dimer): $^{1}H$ NMR (400 MHz, Deuterium Oxide) δ 8.14 (s, 2H), 8.08 (s, 2H), 6.74 (t, J = 1.8 Hz, 2H), (t, J = 6.0 Hz, 2H), 4.59 (t, J = 4.9 Hz, 2H), 4.39 (t, J = 4.8 Hz, 2H), 4.07 (m, 6H). 31P NMR (162 MHz, Deuterium Oxide) δ -12.81. HRMS: calc for [$C_{23}H_{28}N_{13}O_{12}P_2^-$] 740.1461, found 740.1470.

Bis(((2R,3S,4R,5R)-5-(6-amino-9H-purin-9-yl)-3,4-dihydroxytetrahydrofuran-2-yl)methyl) (2-amino-1H-imidazole-3-ium-1,3-diyl)bis(phosphonate) (2AIparaA-imidazolium bridged dimer): $^{1}H$ NMR (400 MHz, Deuterium Oxide) δ 8.11 (s, 2H), 8.10 (s, 2H), 6.65 (t, J = 2.0 Hz, 2H), 6.29 (t, J = 6.5 Hz, 2H), 4.59 (dt, J = 6.3, 3.8 Hz, 2H), 4.14 – 4.05 (m, 4H), 4.02 – 3.94 (m, 2H), 2.70 (dt, J = 13.4, 6.5 Hz, 2H), 2.53 (ddd, J = 14.0, 6.6, 4.4 Hz, 2H). $^{31}P$ NMR (162 MHz, Deuterium Oxide) δ -12.82. HRMS: calc for [$C_{23}H_{28}N_{13}O_{12}P_2^-$] 740.1461, found 740.1474.

Bis(((2S,4R,5R)-5-(6-amino-9H-purin-9-yl)-4-hydroxytetrahydrofuran-2-yl)methyl) (2-amino-1H-imidazole-3-ium-1,3-diyl)bis(phosphonate) (2AIpdA-imidazolium bridged dimer): $^{1}H$ NMR (400 MHz, Deuterium Oxide) δ 8.11 (s, 2H), 8.10 (s, 2H), 6.65 (t, J = 2.0 Hz, 2H), 6.29 (t, J = 6.5 Hz, 2H), 4.59 (dt, J = 6.3, 3.8 Hz, 2H), 4.14 – 4.05 (m, 4H), 4.02 – 3.94 (m, 2H), 2.70 (dt, J = 13.4, 6.5 Hz, 2H), 2.53 (ddd, J = 14.0, 6.6, 4.4 Hz, 2H). 31P NMR (162 MHz, Deuterium Oxide) δ -12.82. HRMS: calc for [$C_{23}H_{28}N_{13}O_{10}P_2^-$] 708.1563, found 708.1554.

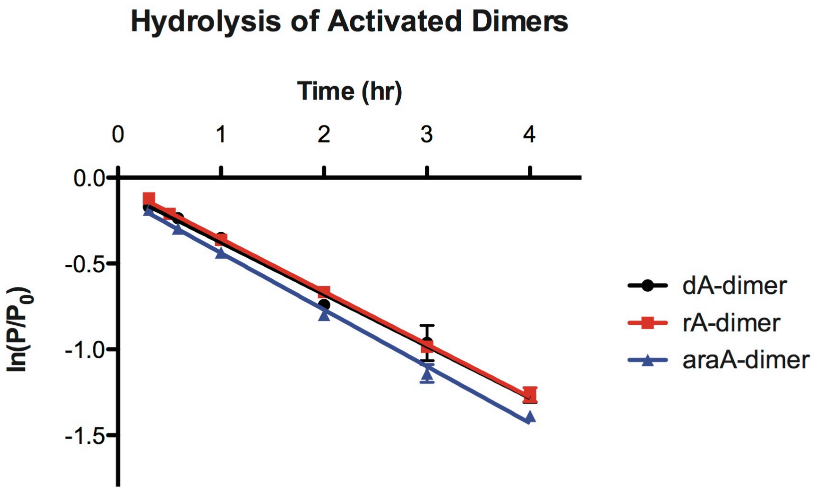


$k_{dA} = 0.30 \pm 0.01\ hr^{-1}$

$k_{rA} = 0.31 \pm 0.01\ hr^{-1}$

$k_{araA} = 0.33 \pm 0.01\ hr^{-1}$

**Figure S1.** Kinetic analysis of the hydrolysis of imidazolium bridged dimers for dA, rA, and araA was carried out in triplicate using 5 mM imidazolium bridged dimers, 50 mM $MgCl_2$, 200 mM $Na^+$-HEPES pH 8.0. As standard for first-order kinetics, a linearized plot of $\ln(P/P_0)$ as a function of time is shown above and the rate of extension was determined from linear least-squares fits of the data from an average of three independent experiments.

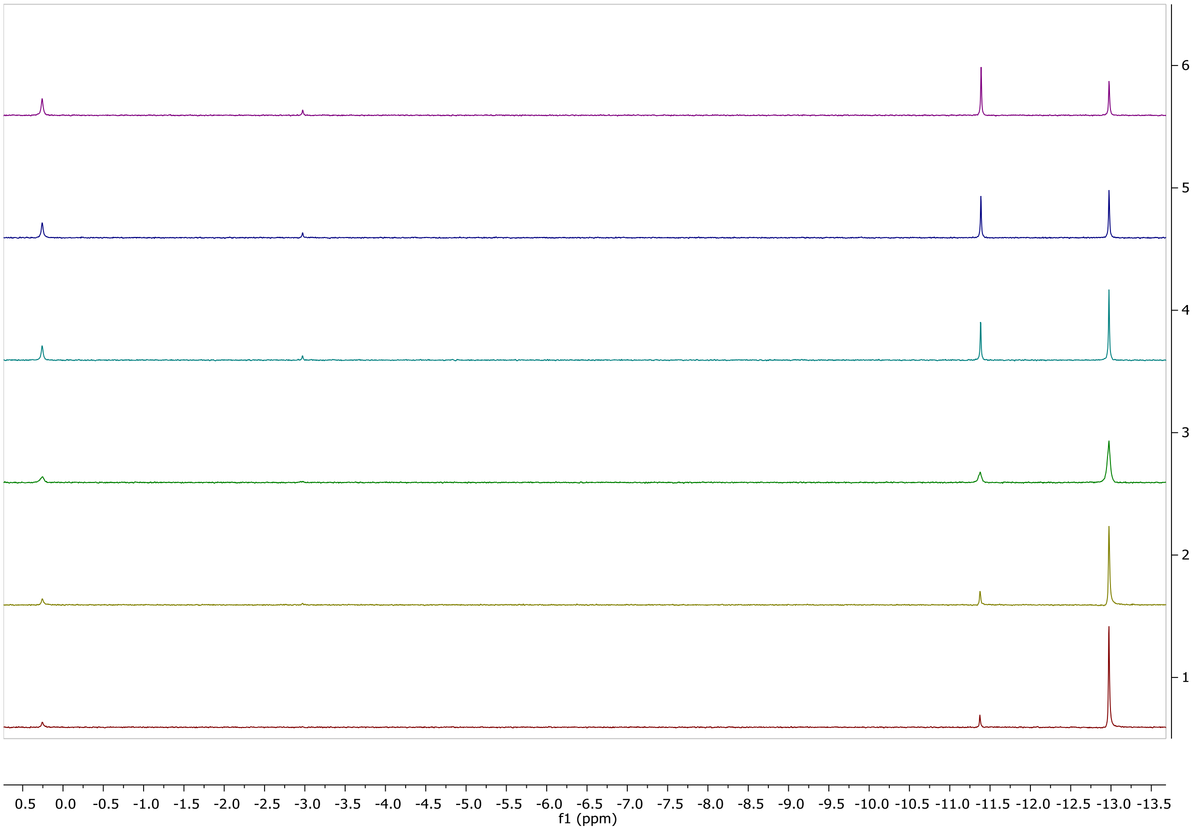


**Figure S2.** [31]P NMR spectrum of the hydrolysis of dA imidazolium bridged dimer under nonenzymatic primer extension conditions (5 mM imidazolium bridged dimer, 50 mM $MgCl_2$, 200 mM $Na^+$-HEPES pH 8.0, 10% $D_2O$) after 18, 35, 60, 120, 180, and 240 minutes of reaction time (bottom to top).

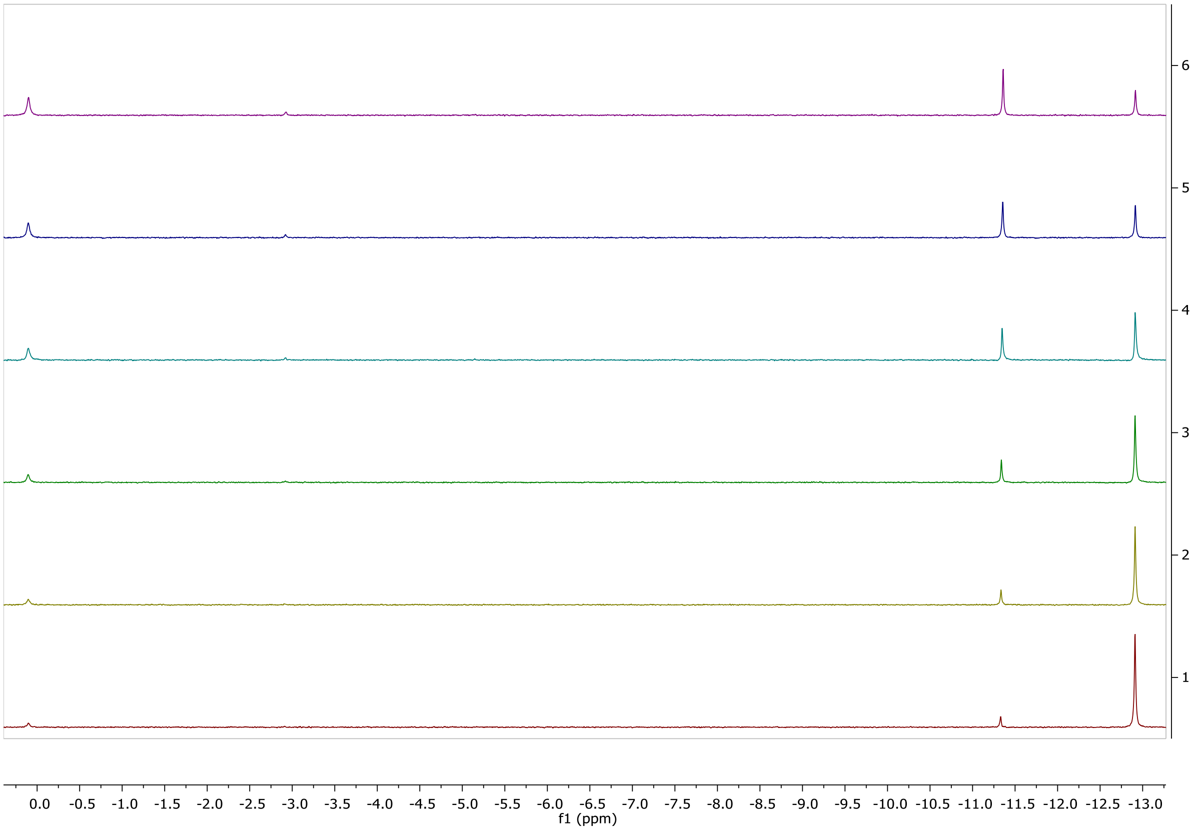


**Figure S3.** $^{31}P$ NMR spectrum of the hydrolysis of araA imidazolium bridged dimer under nonenzymatic primer extension conditions (5 mM imidazolium bridged dimer, 50 mM $MgCl_2$, 200 mM $Na^+$-HEPES pH 8.0, 10% $D_2O$) after 18, 35, 60, 120, 180, and 240 minutes of reaction time (bottom to top).

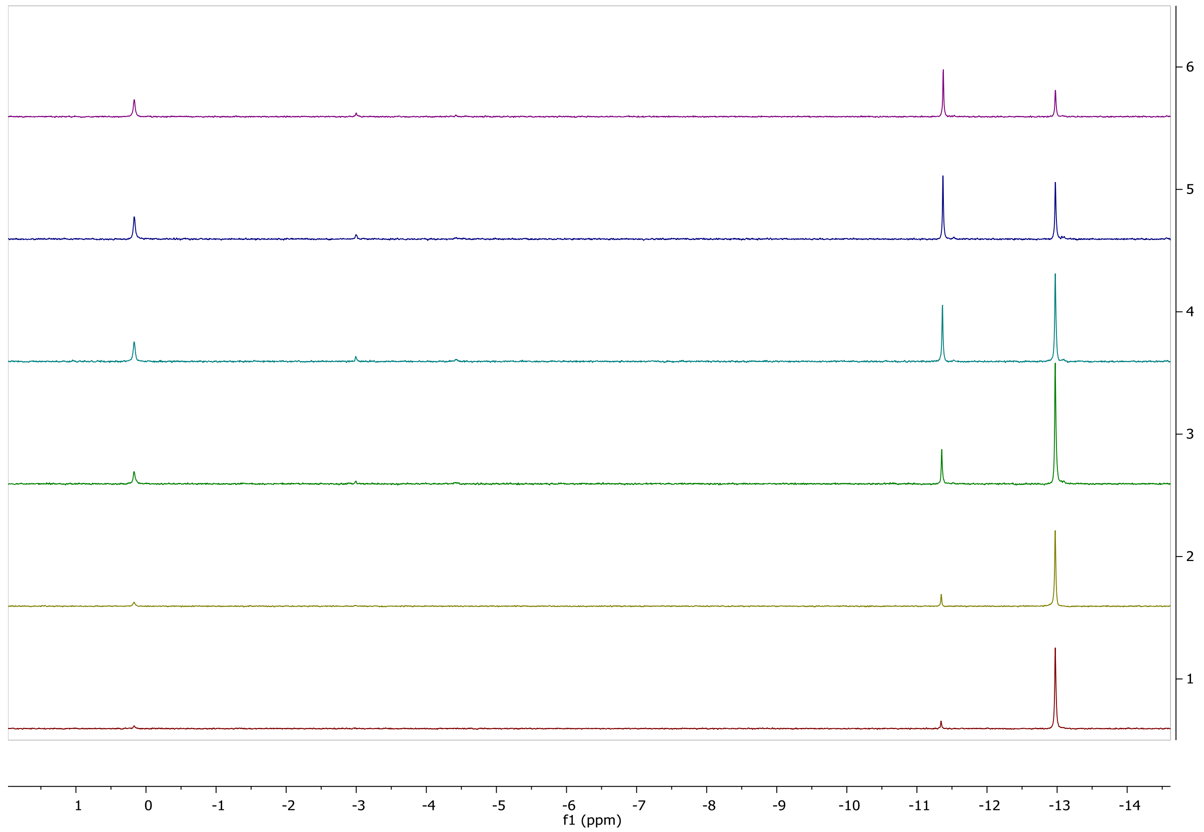


**Figure S4.** $^{31}P$ NMR spectrum of the hydrolysis of rA imidazolium bridged dimer under nonenzymatic primer extension conditions (5 mM imidazolium bridged dimer, 50 mM $MgCl_2$, 200 mM $Na^+$-HEPES pH 8.0, 10% $D_2O$) after 18, 30, 60, 120, 180, and 240 minutes of reaction time (bottom to top).

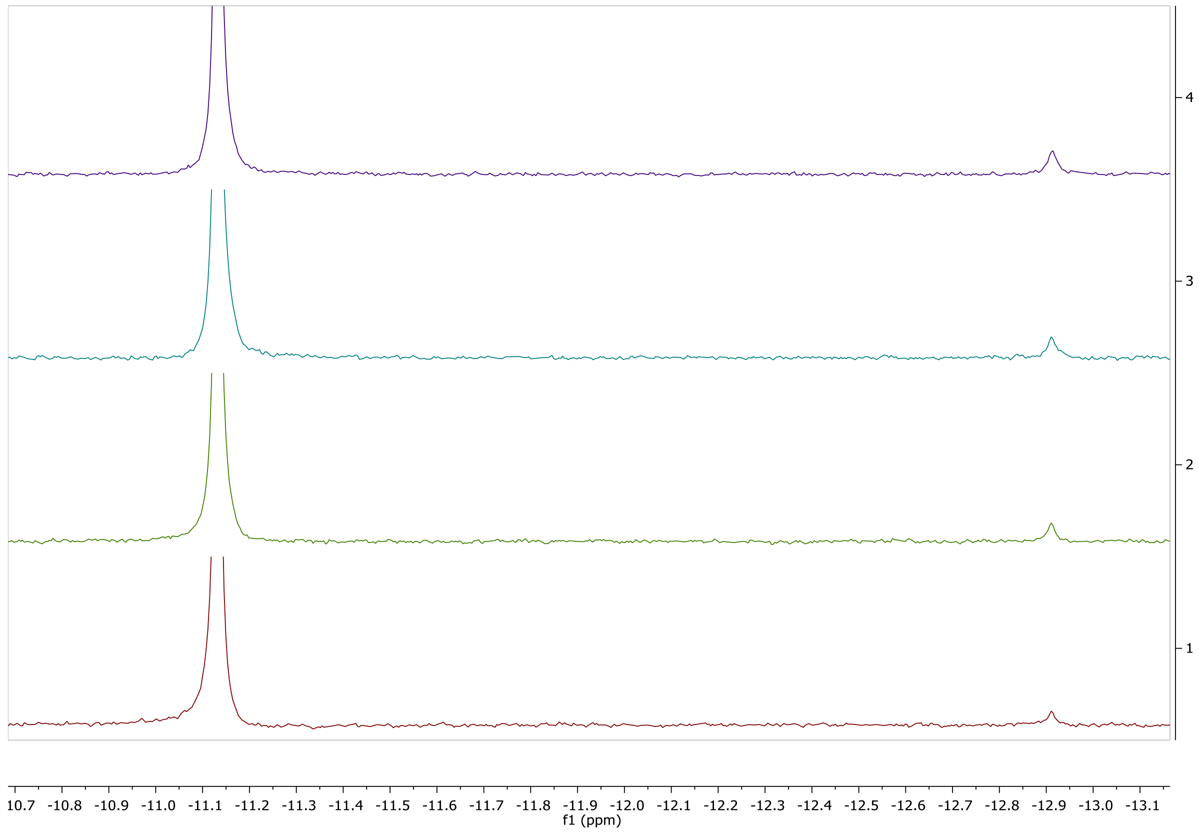


**Figure S5.** $^{31}P$ NMR spectrum of the formation of araA imidazolium bridged dimer under nonenzymatic primer extension conditions (24 mM 2AIparaA, 100 mM $MgCl_2$, 100 mM $Na^+$-HEPES pH 8.0, 10% $D_2O$) after 7, 16, 25, and 36 minutes of reaction time (bottom to top). Kinetic analysis of the hydrolysis of imidazolium bridged dimers was determined to be 3.1(3) x $10^{-3}$ $hr^{-1}$ $mM^{-1}$.

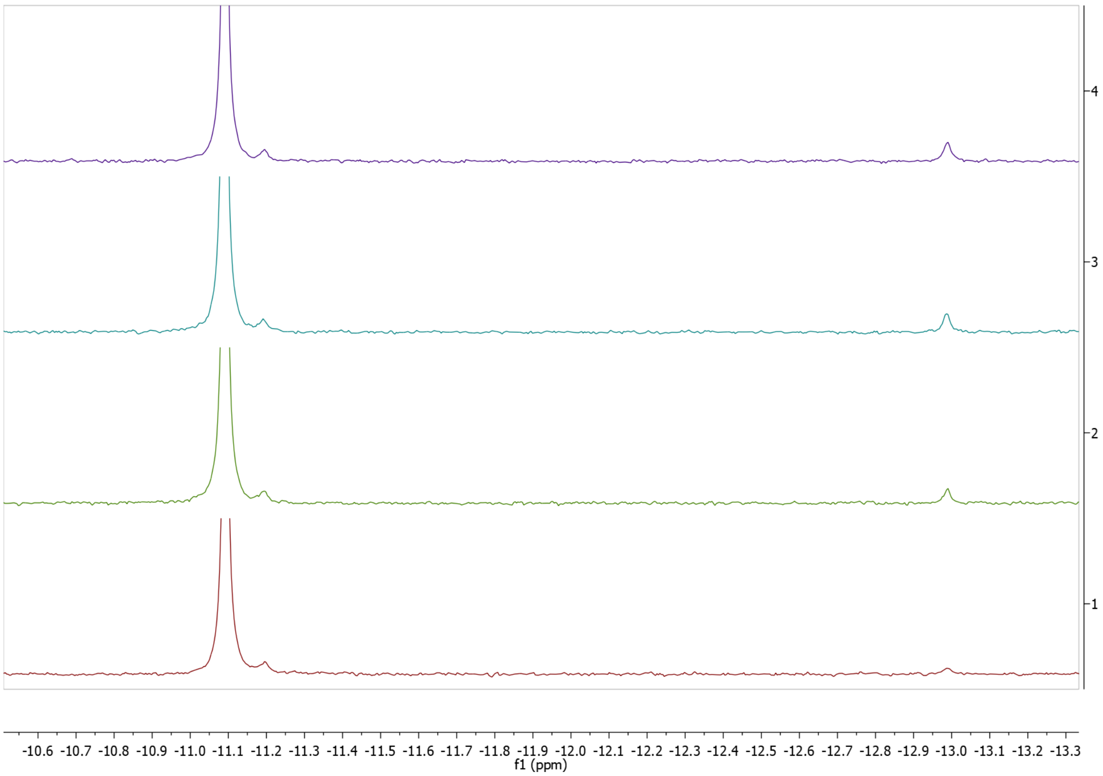


**Figure S6.** $^{31}P$ NMR spectrum of the formation of rA imidazolium bridged dimer under nonenzymatic primer extension conditions (24 mM 2AIprA, 100 mM $MgCl_2$, 100 mM $Na^+$-HEPES pH 8.0, 10% $D_2O$) after 7, 16, 25, and 36 minutes of reaction time (bottom to top). Kinetic analysis of the hydrolysis of imidazolium bridged dimers was determined to be 3.2(2) x $10^{-3}$ $hr^{-1}$ $mM^{-1}$.

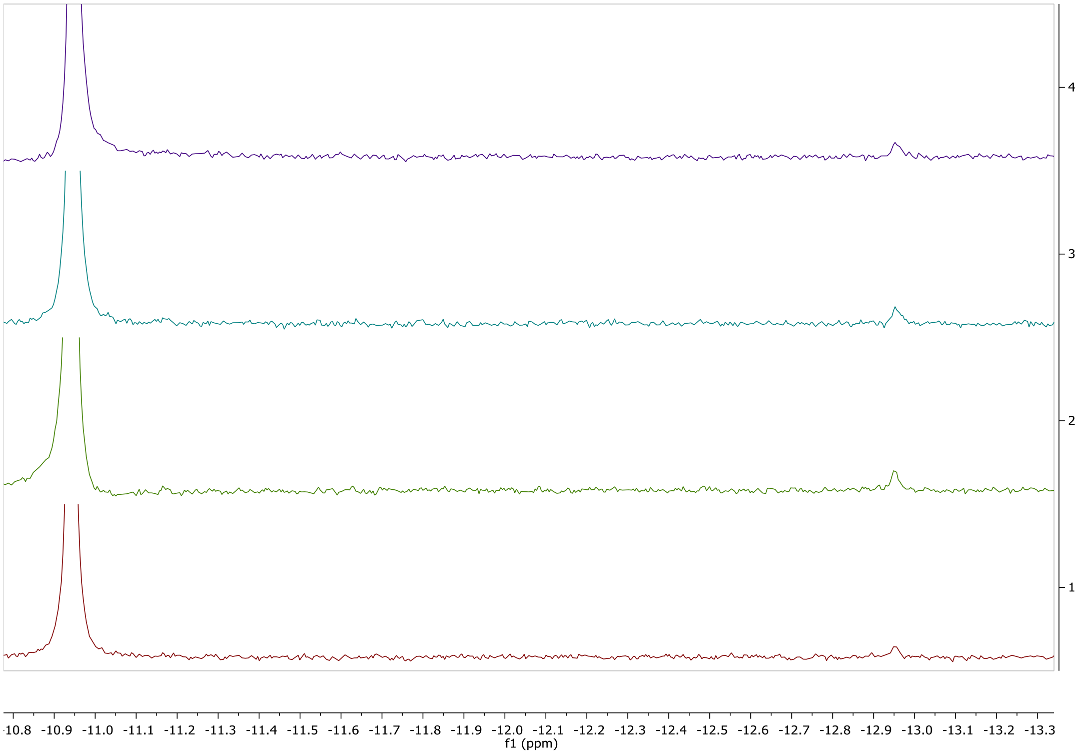


**Figure S7.** $^{31}P$ NMR spectrum of the formation of dA imidazolium bridged dimer under nonenzymatic primer extension conditions (24 mM 2AIpdA, 100 mM $MgCl_2$, 100 mM $Na^+$-HEPES pH 8.0, 10% $D_2O$) after 7, 16, 25, and 36 minutes of reaction time (bottom to top). Kinetic analysis of the hydrolysis of imidazolium bridged dimers was determined to be 2.8(2) x $10^{-3}$ $hr^{-1}$ $mM^{-1}$.

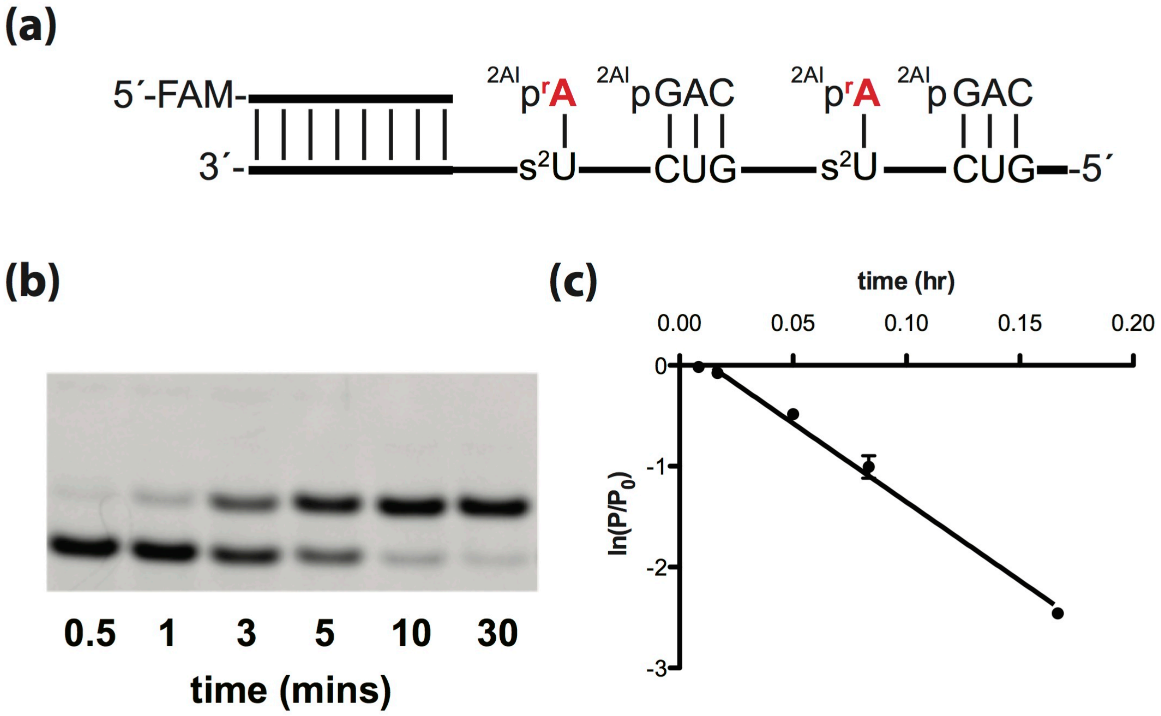


$k = 16 \pm 1\ \text{hr}^{-1}$

**Figure S8.** Primer extension reactions were carried out in triplicate using 20 mM 2AIprA, 0.5 mM 2AIpGAC, 50 mM $MgCl_2$, 200 mM $Na^+$-HEPES pH 8.0. (a) Schematic representation of a primer extension reaction. (b) Representative PAGE analysis of result. (c) Plot of $\ln(P/P_0)$ as a function of time. The rate of extension was determined from linear least-squares fits of the data from three independent experiments.

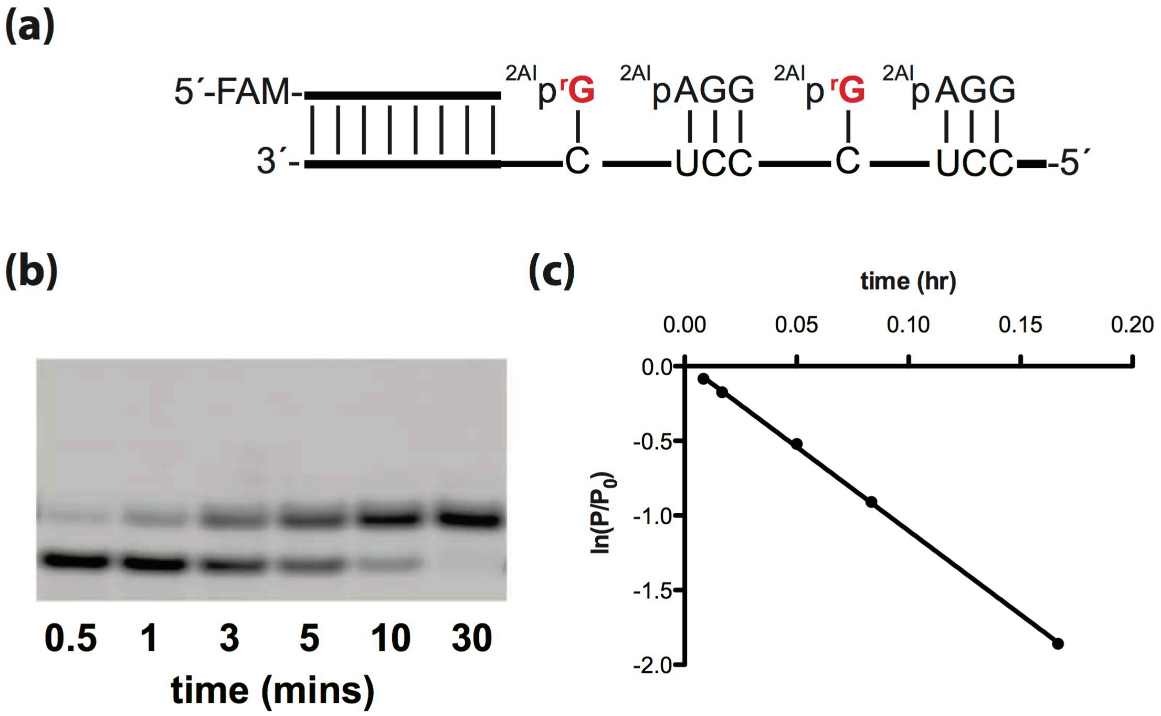


***k*= 11 ± 1 hr$^{-1}$**

**Figure S9.** Primer extension reactions were carried out in triplicate using 20 mM 2AIprG, 0.5 mM 2AIpGAC, 50 mM $MgCl_2$, 200 mM $Na^+$-HEPES pH 8.0. (a) Schematic representation of a primer extension reaction. (b) Representative PAGE analysis of result. (c) Plot of $\ln(P/P_0)$ as a function of time. The rate of extension was determined from linear least-squares fits of the data from three independent experiments.

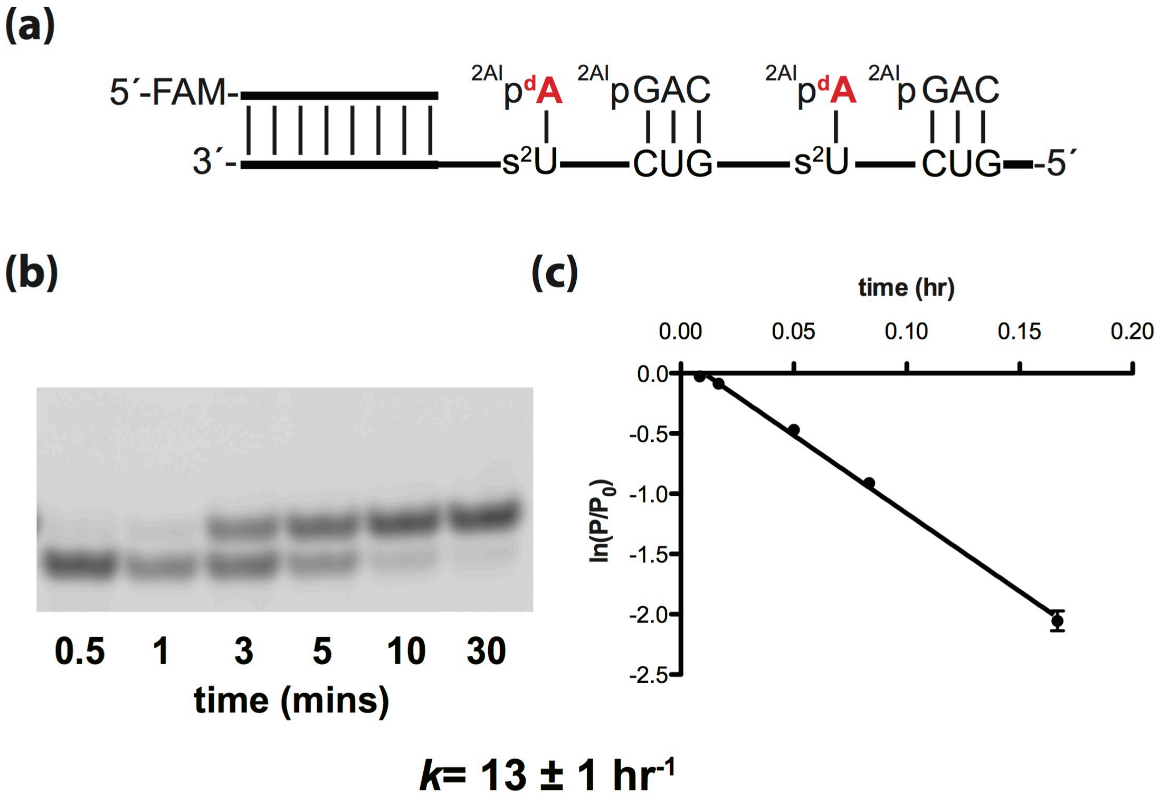


**Figure S10.** Primer extension reactions were carried out in triplicate using 20 mM 2AIpdA, 0.5 mM 2AIpGAC, 50 mM $MgCl_2$, 200 mM $Na^+$-HEPES pH 8.0. (a) Schematic representation of a primer extension reaction. (b) Representative PAGE analysis of result. (c) Plot of $\ln(P/P_0)$ as a function of time. The rate of extension was determined from linear least-squares fits of the data from three independent experiments.

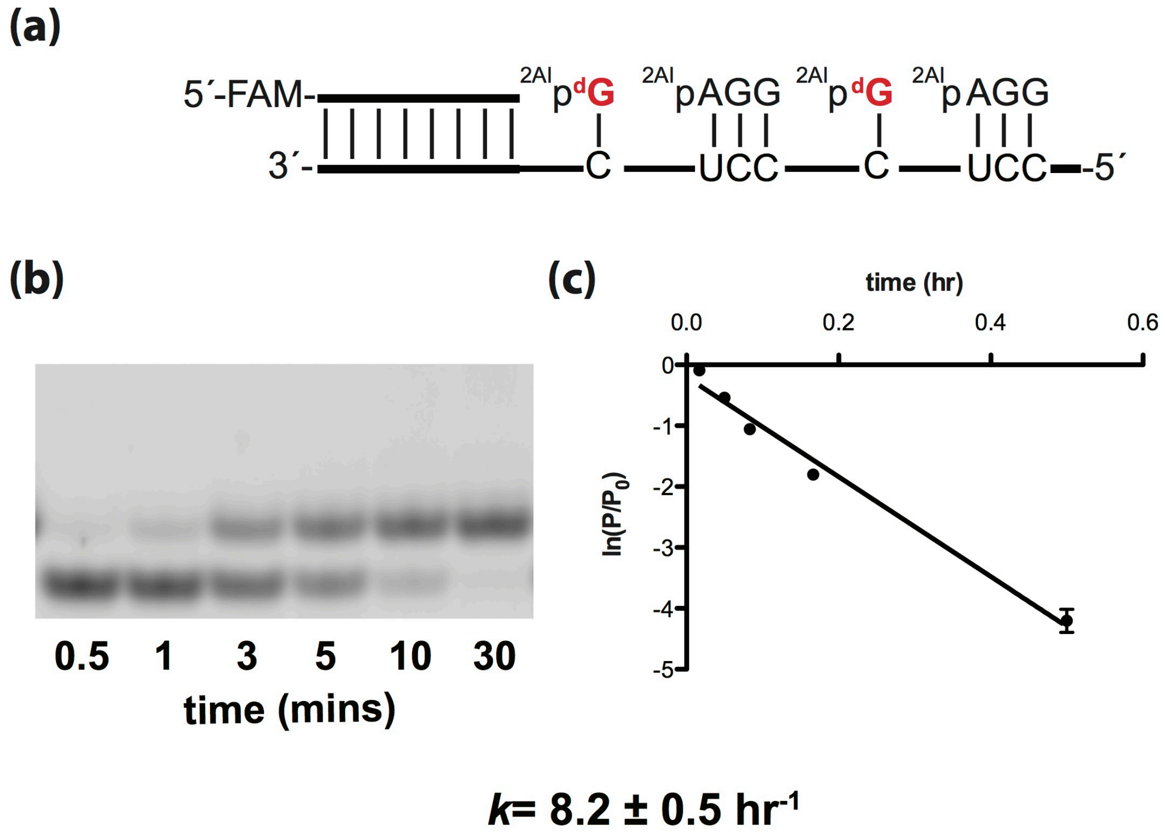


**Figure S11.** Primer extension reactions were carried out in triplicate using 20 mM 2AIpdG, 0.5 mM 2AIpGAC, 50 mM $MgCl_2$, 200 mM $Na^+$-HEPES pH 8.0. (a) Schematic representation of a primer extension reaction. (b) Representative PAGE analysis of result. (c) Plot of $\ln(P/P_0)$ as a function of time. The rate of extension was determined from linear least-squares fits of the data from three independent experiments.

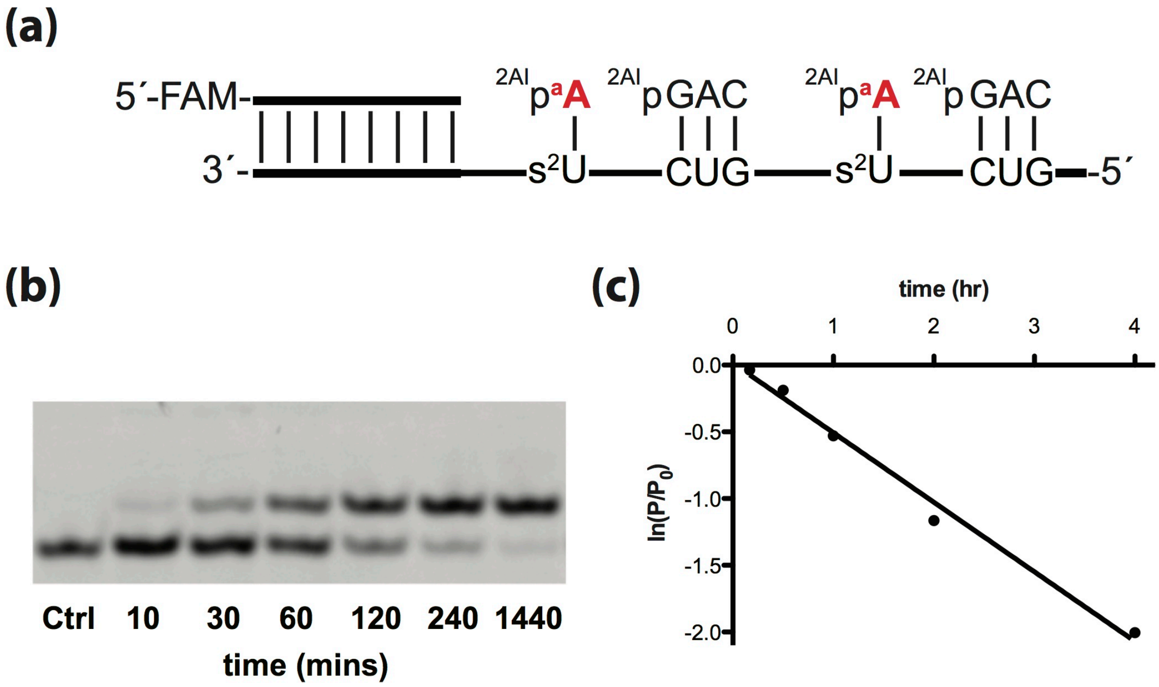


$k = 0.52 \pm 0.02\ hr^{-1}$

**Figure S12.** Primer extension reactions were carried out in triplicate using 20 mM 2AIparaA, 0.5 mM 2AIpGAC, 50 mM $MgCl_2$, 200 mM $Na^+$-HEPES pH 8.0. (a) Schematic representation of a primer extension reaction. (b) Representative PAGE analysis of result. (c) Plot of $\ln(P/P_0)$ as a function of time. The rate of extension was determined from linear least-squares fits of the data from three independent experiments.

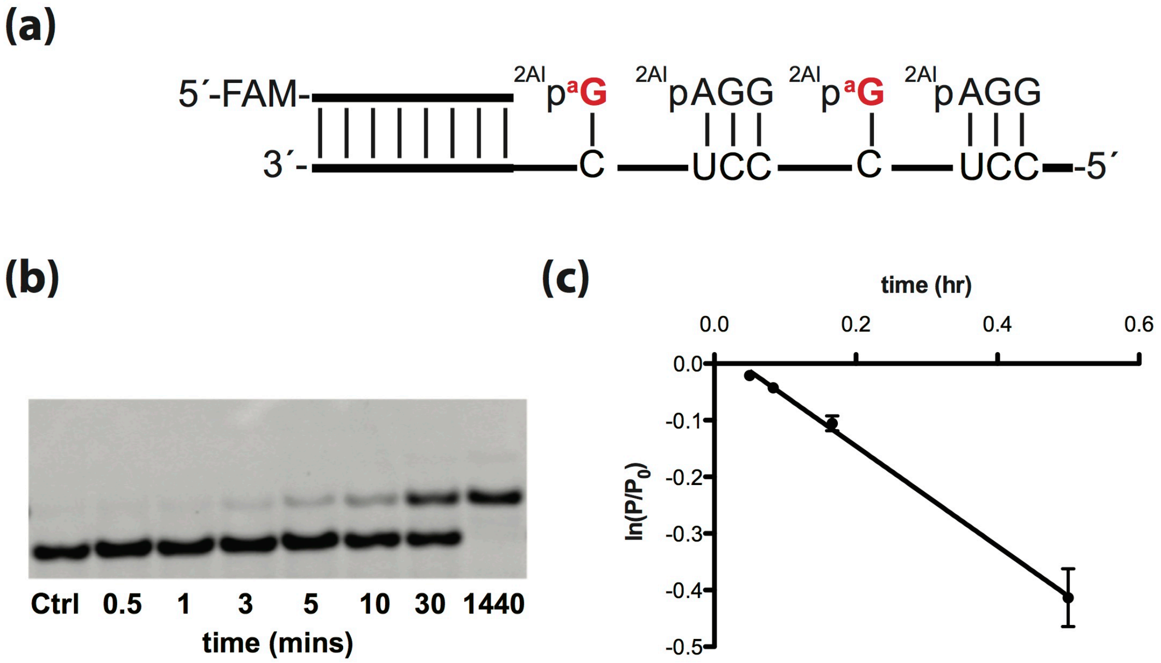


$k$= 0.88 ± 0.04 hr$^{-1}$

**Figure S13.** Primer extension reactions were carried out in triplicate using 20 mM 2AIparaG, 0.5 mM 2AIpGAC, 50 mM $MgCl_2$, 200 mM $Na^+$-HEPES pH 8.0. (a) Schematic representation of a primer extension reaction. (b) Representative PAGE analysis of result. (c) Plot of ln($P/P_0$) as a function of time. The rate of extension was determined from linear least-squares fits of the data from three independent experiments.

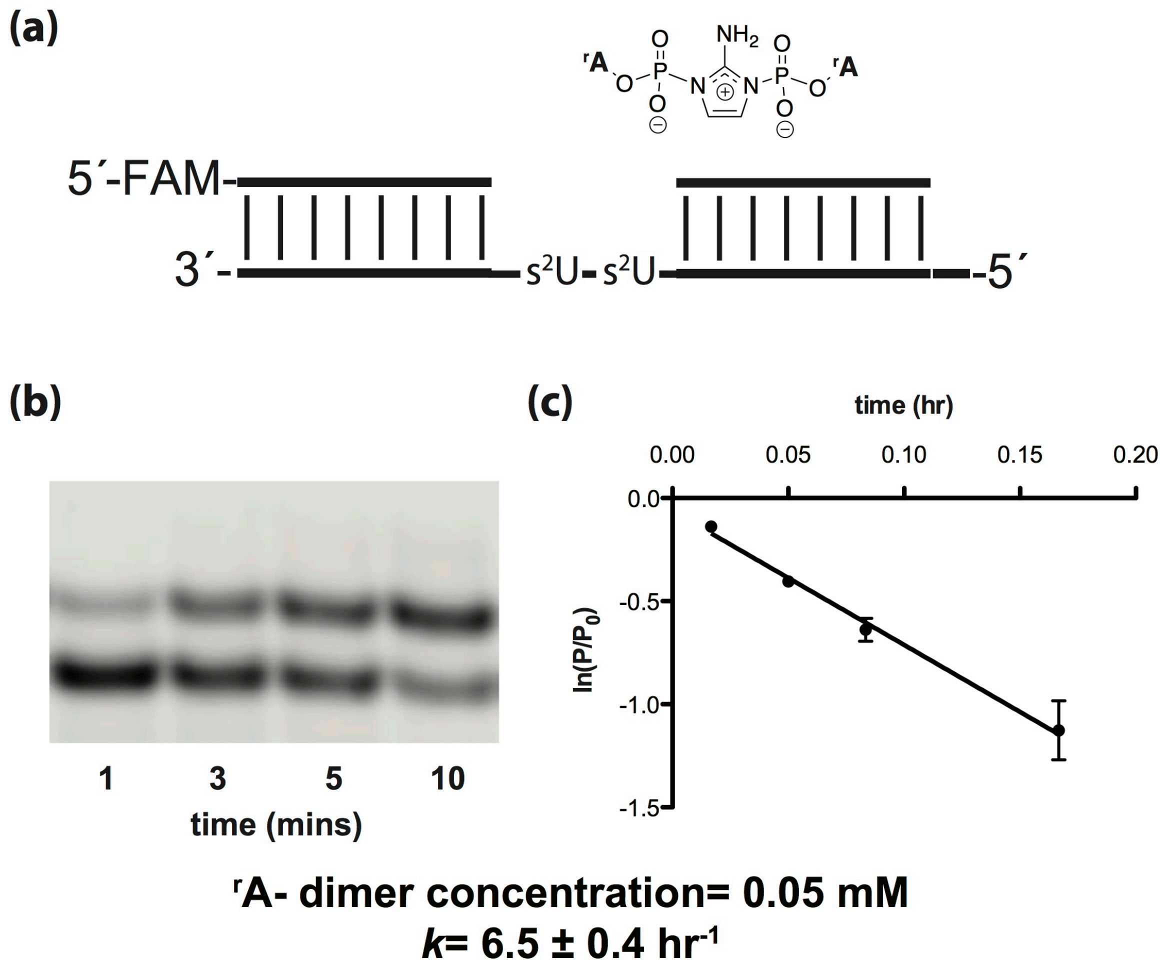


**Figure S14.** Michaelis-Menten kinetics of the addition of 2AIprA imidazolium-bridged dimer. Primer extension reactions were carried out in triplicate using 0.05 mM of 2AIpA imidazolium-bridged dimer, 50 mM $MgCl_2$, 200 mM $Na^+$-HEPES pH 8.0. (a) Schematic representation of a primer extension reaction. (b) Representative PAGE analysis of result. (c) Plot of $\ln(P/P_0)$as a function of time. The rate of extension was determined from linear least-squares fits of the data from three independent experiments.

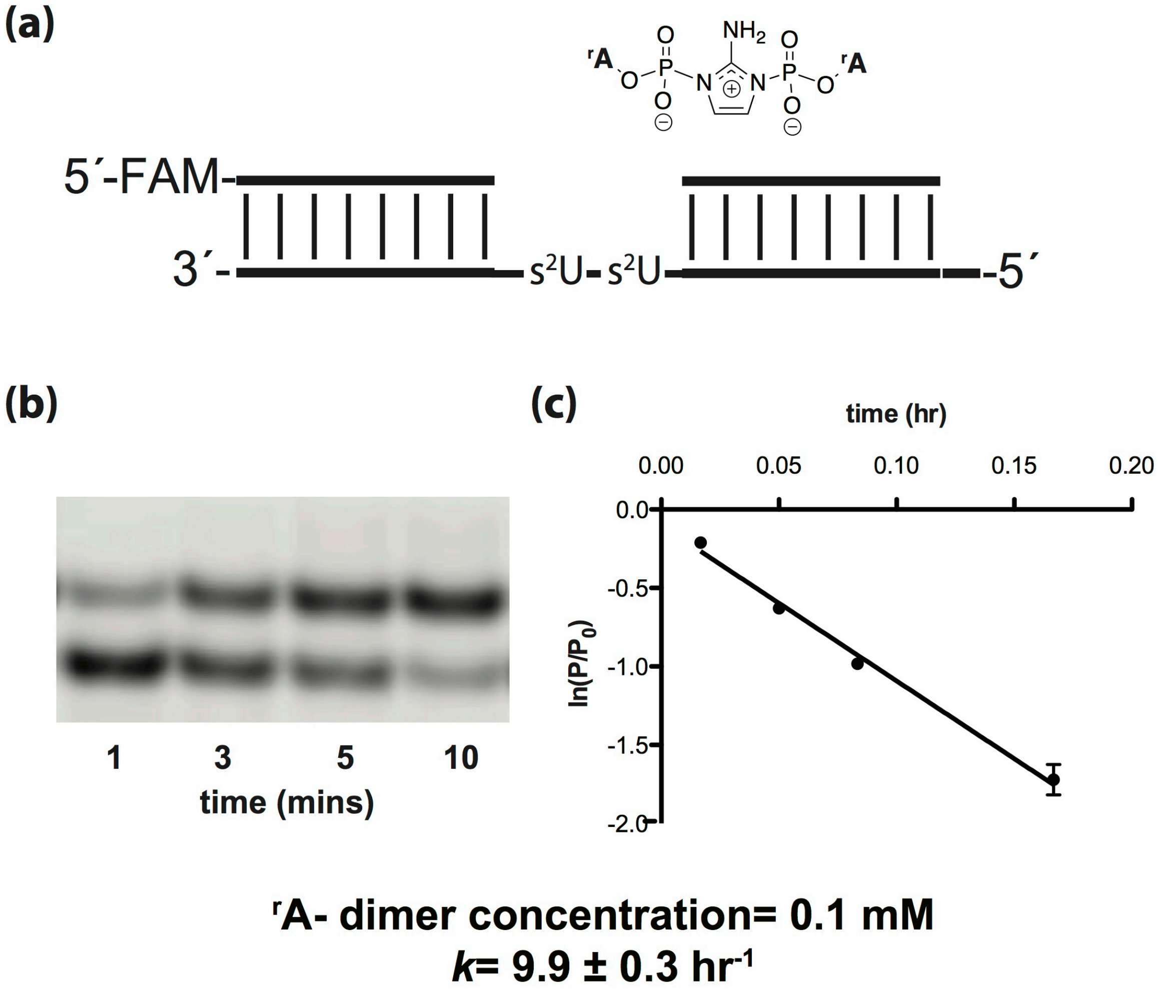


**Figure S15.** Michaelis-Menten kinetics of the addition of 2AIprA imidazolium-bridged dimer. Primer extension reactions were carried out in triplicate using 0.1 mM of 2AIpA imidazolium-bridged dimer, 50 mM $MgCl_2$, 200 mM $Na^+$-HEPES pH 8.0. (a) Schematic representation of a primer extension reaction. (b) Representative PAGE analysis of result. (c) Plot of $\ln(P/P_0)$as a function of time. The rate of extension was determined from linear least-squares fits of the data from three independent experiments.

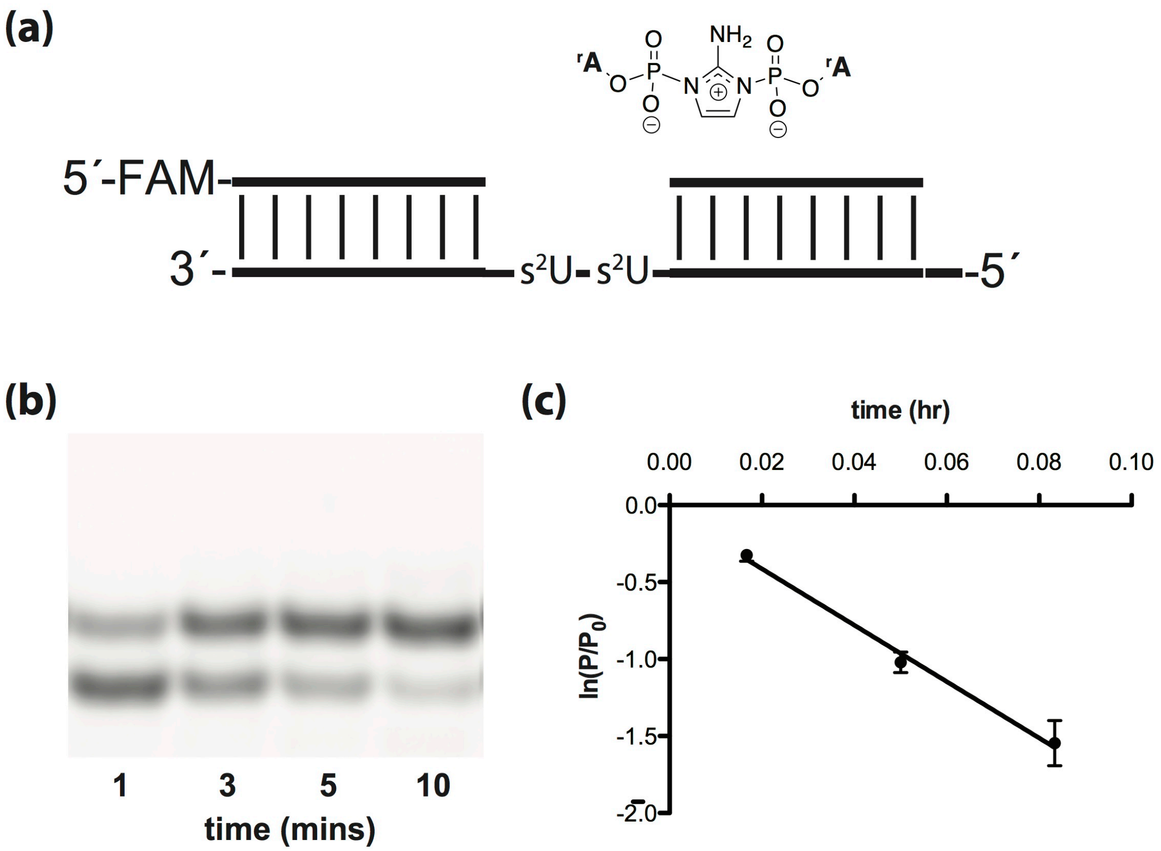


**$^r$A- dimer concentration= 0.5 mM**
***k*= 18 ± 1 hr$^{-1}$**

**Figure S16.** Michaelis-Menten kinetics of the addition of 2AIprA imidazolium-bridged dimer. Primer extension reactions were carried out in triplicate using 0.5 mM of 2AIpA imidazolium-bridged dimer, 50 mM $MgCl_2$, 200 mM $Na^+$-HEPES pH 8.0. (a) Schematic representation of a primer extension reaction. (b) Representative PAGE analysis of result. (c) Plot of $\ln(P/P_0)$as a function of time. The rate of extension was determined from linear least-squares fits of the data from three independent experiments.

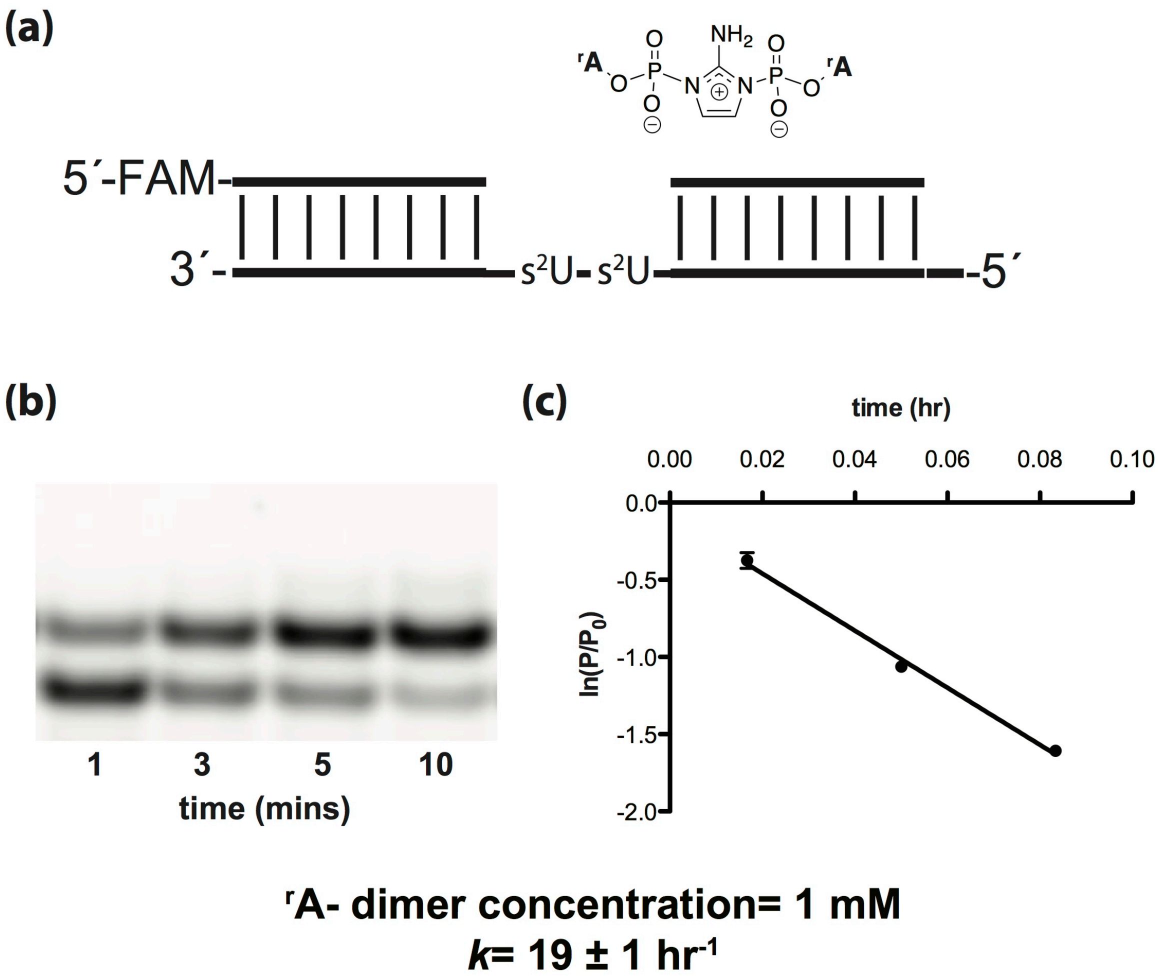


**Figure S17.** Michaelis-Menten kinetics of the addition of 2AIprA imidazolium-bridged dimer. Primer extension reactions were carried out in triplicate using 1 mM of 2AIpA imidazolium-bridged dimer, 50 mM $MgCl_2$, 200 mM $Na^+$-HEPES pH 8.0. (a) Schematic representation of a primer extension reaction. (b) Representative PAGE analysis of result. (c) Plot of $\ln(P/P_0)$as a function of time. The rate of extension was determined from linear least-squares fits of the data from three independent experiments.

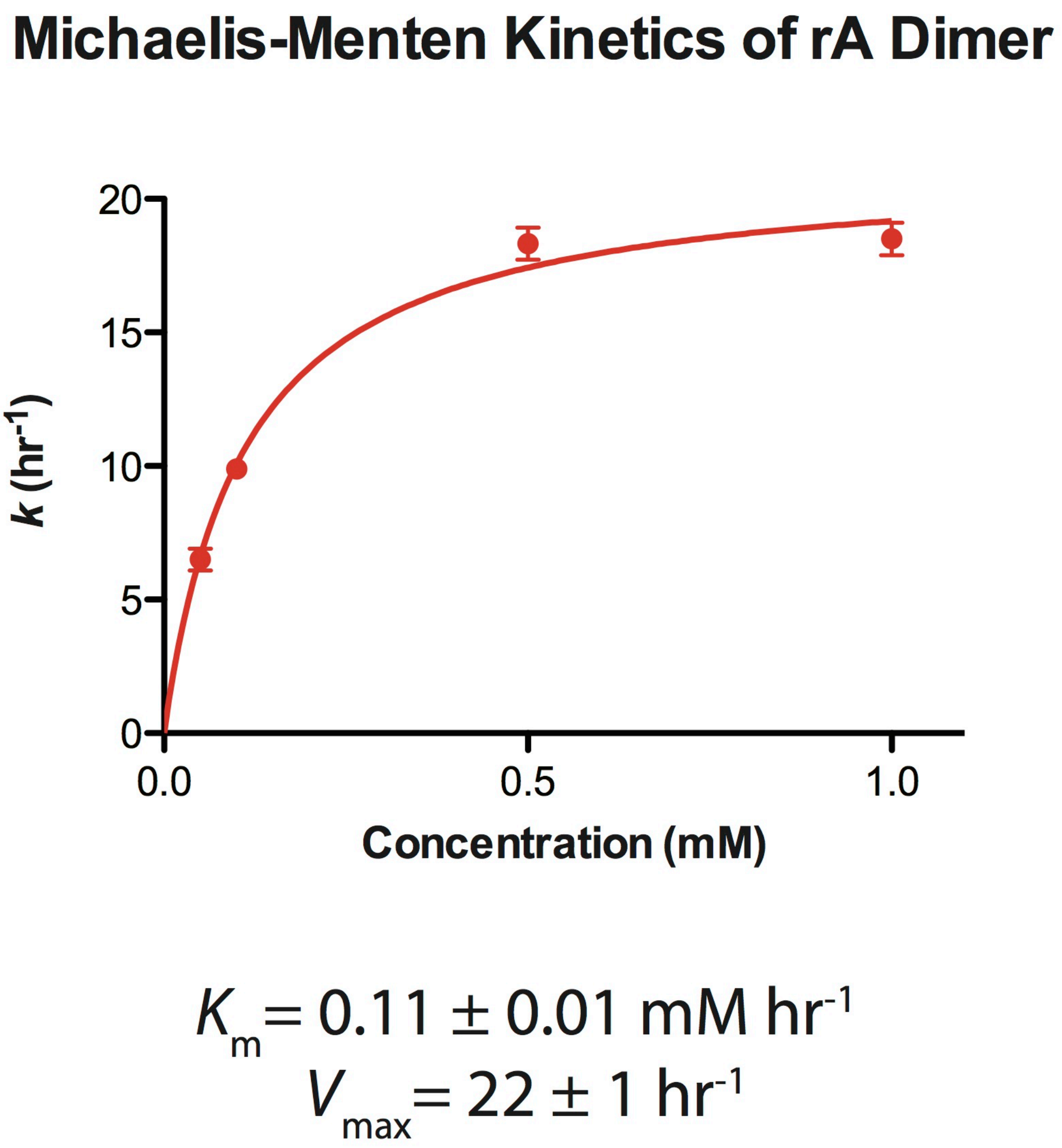


**Figure S18.** Michaelis-Menten kinetics of the addition of 2AIprA imidazolium-bridged dimer. Primer extension reactions were carried out in triplicate using various concentrations of 2AIpA imidazolium-bridged dimer, 50 mM $MgCl_2$, 200 mM $Na^+$-HEPES pH 8.0 and plot of *k* ($hr^{-1}$) as a function of the concentration (mM) of 2AIprA imidazolium-bridged dimer is shown as above. Michaelis-Menten parameters are $K_M$ : 0.11(1) mM and $V_{MAX}$ of 21.3(5) mM $hr^{-1}$.

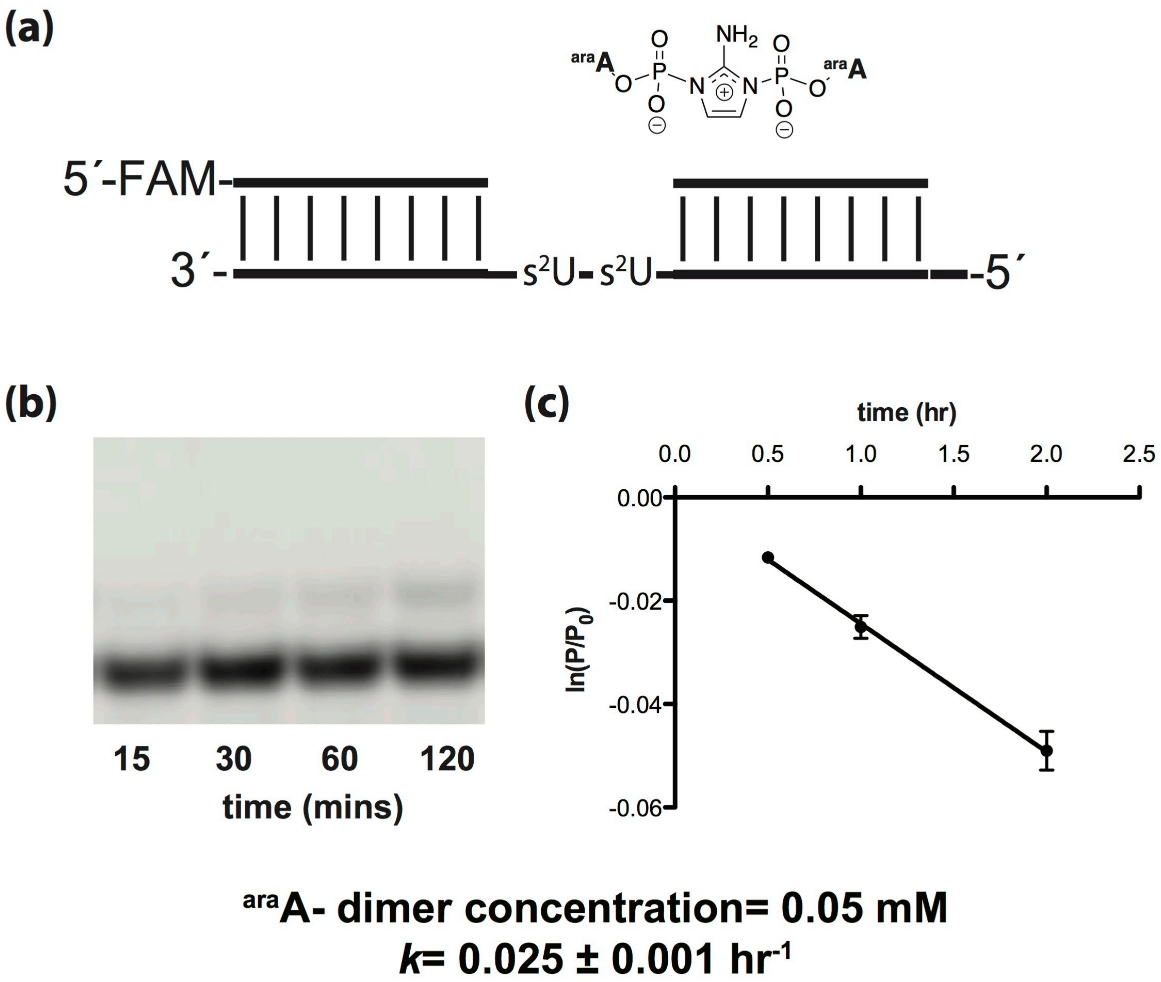


**Figure S19.** Michaelis-Menten kinetics of the addition of 2AIparaA imidazolium-bridged dimer. Primer extension reactions were carried out in triplicate using 0.05 mM of 2AIparaA imidazolium-bridged dimer, 50 mM $MgCl_2$, 200 mM $Na^+$-HEPES pH 8.0. (a) Schematic representation of a primer extension reaction. (b) Representative PAGE analysis of result. (c) Plot of $\ln(P/P_0)$as a function of time. The rate of extension was determined from linear least-squares fits of the data from three independent experiments.

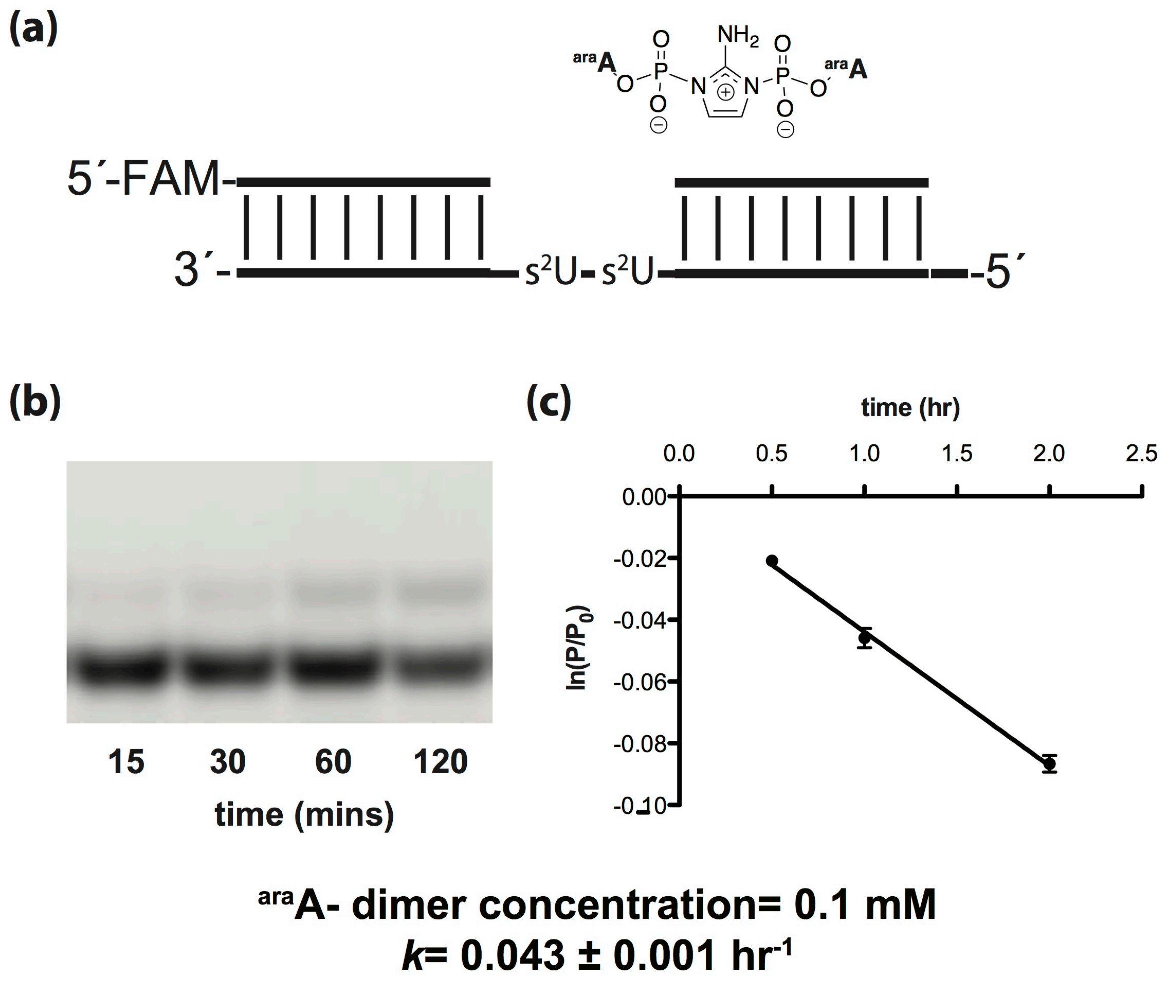


**Figure S20.** Michaelis-Menten kinetics of the addition of 2AIparaA imidazolium-bridged dimer. Primer extension reactions were carried out in triplicate using 0.1 mM of 2AIparaA imidazolium-bridged dimer, 50 mM $MgCl_2$, 200 mM $Na^+$-HEPES pH 8.0. (a) Schematic representation of a primer extension reaction. (b) Representative PAGE analysis of result. (c) Plot of $\ln(P/P_0)$as a function of time. The rate of extension was determined from linear least-squares fits of the data from three independent experiments.

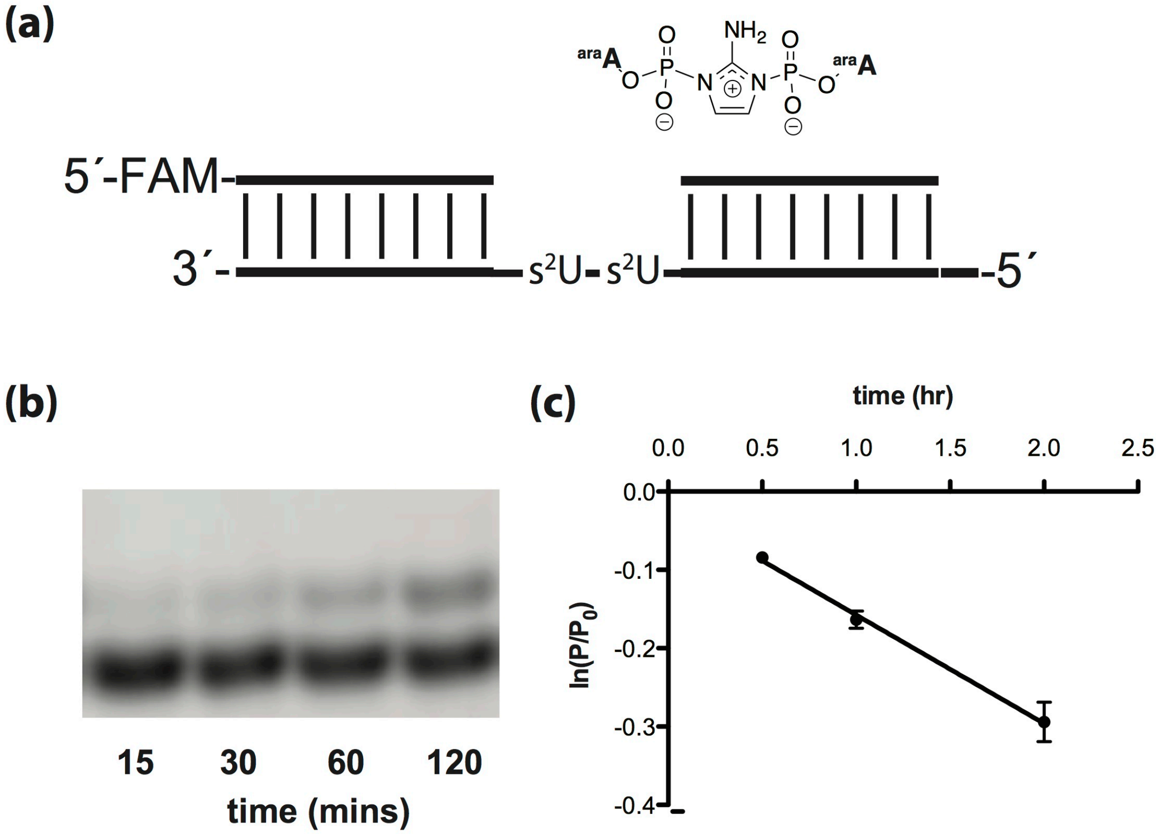


**$^{ara}$A- dimer concentration= 0.5 mM**

**$k$= 0.14 ± 0.01 hr$^{-1}$**

**Figure S21.** Michaelis-Menten kinetics of the addition of 2AIparaA imidazolium-bridged dimer. Primer extension reactions were carried out in triplicate using 0.5 mM of 2AIparaA imidazolium-bridged dimer, 50 mM $MgCl_2$, 200 mM $Na^+$-HEPES pH 8.0. (a) Schematic representation of a primer extension reaction. (b) Representative PAGE analysis of result. (c) Plot of ln($P/P_0$)as a function of time. The rate of extension was determined from linear least-squares fits of the data from three independent experiments.

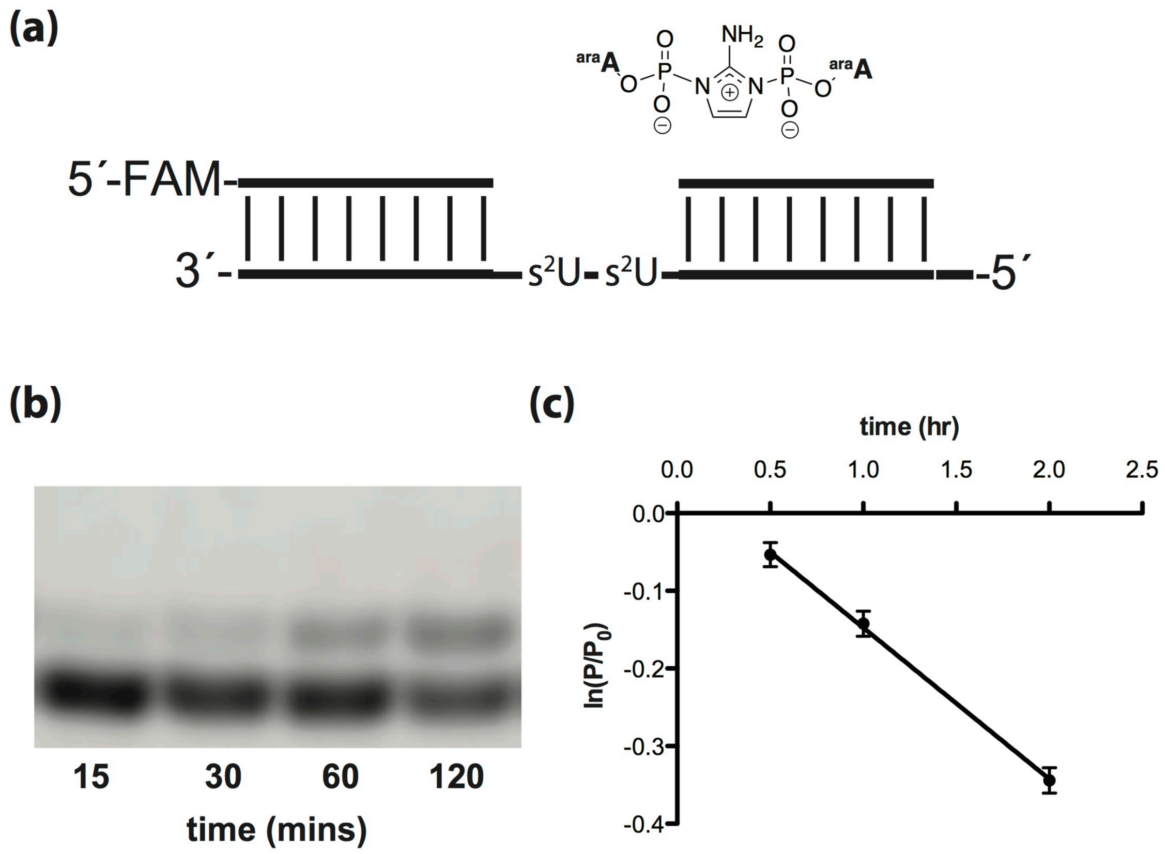


**$^{ara}$A- dimer concentration= 1 mM**
**$k$= 0.20 ± 0.01 hr$^{-1}$**

**Figure S22.** Michaelis-Menten kinetics of the addition of 2AIparaA imidazolium-bridged dimer. Primer extension reactions were carried out in triplicate using 1 mM of 2AIparaA imidazolium-bridged dimer, 50 mM $MgCl_2$, 200 mM $Na^+$-HEPES pH 8.0. (a) Schematic representation of a primer extension reaction. (b) Representative PAGE analysis of result. (c) Plot of $\ln(P/P_0)$as a function of time. The rate of extension was determined from linear least-squares fits of the data from three independent experiments.

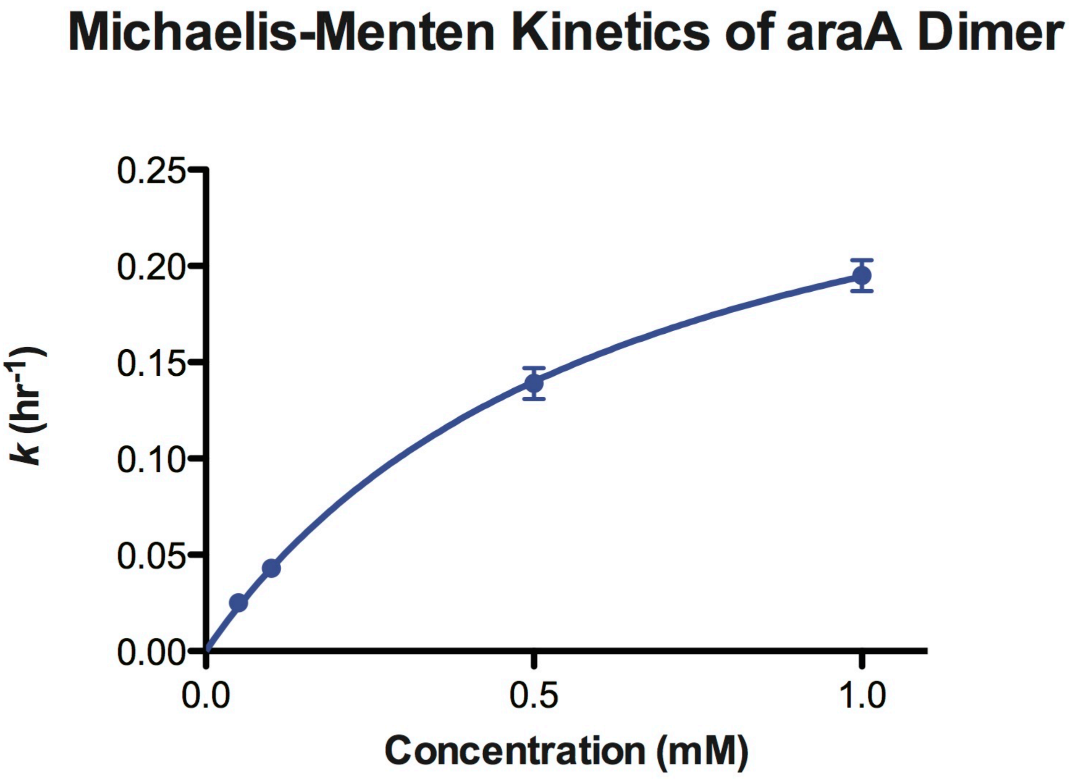


$$K_m = 0.64 \pm 0.07 \text{ mM hr}^{-1}$$
$$V_{max} = 0.32 \pm 0.02 \text{ hr}^{-1}$$

**Figure S23.** Michaelis-Menten kinetics of the addition of 2AIparaA imidazolium-bridged dimer. Primer extension reactions were carried out in triplicate using various concentrations of 2AIparaA imidazolium-bridged dimer, 50 mM $MgCl_2$, 200 mM $Na^+$-HEPES pH 8.0 and plot of *k* ($hr^{-1}$) as a function of the concentration (mM) of 2AIparaA imidazolium-bridged dimer is shown as above. Michaelis-Menten parameters are $K_M$ : 0.64(7) mM and $V_{MAX}$ of 0.32(2) mM $hr^{-1}$.

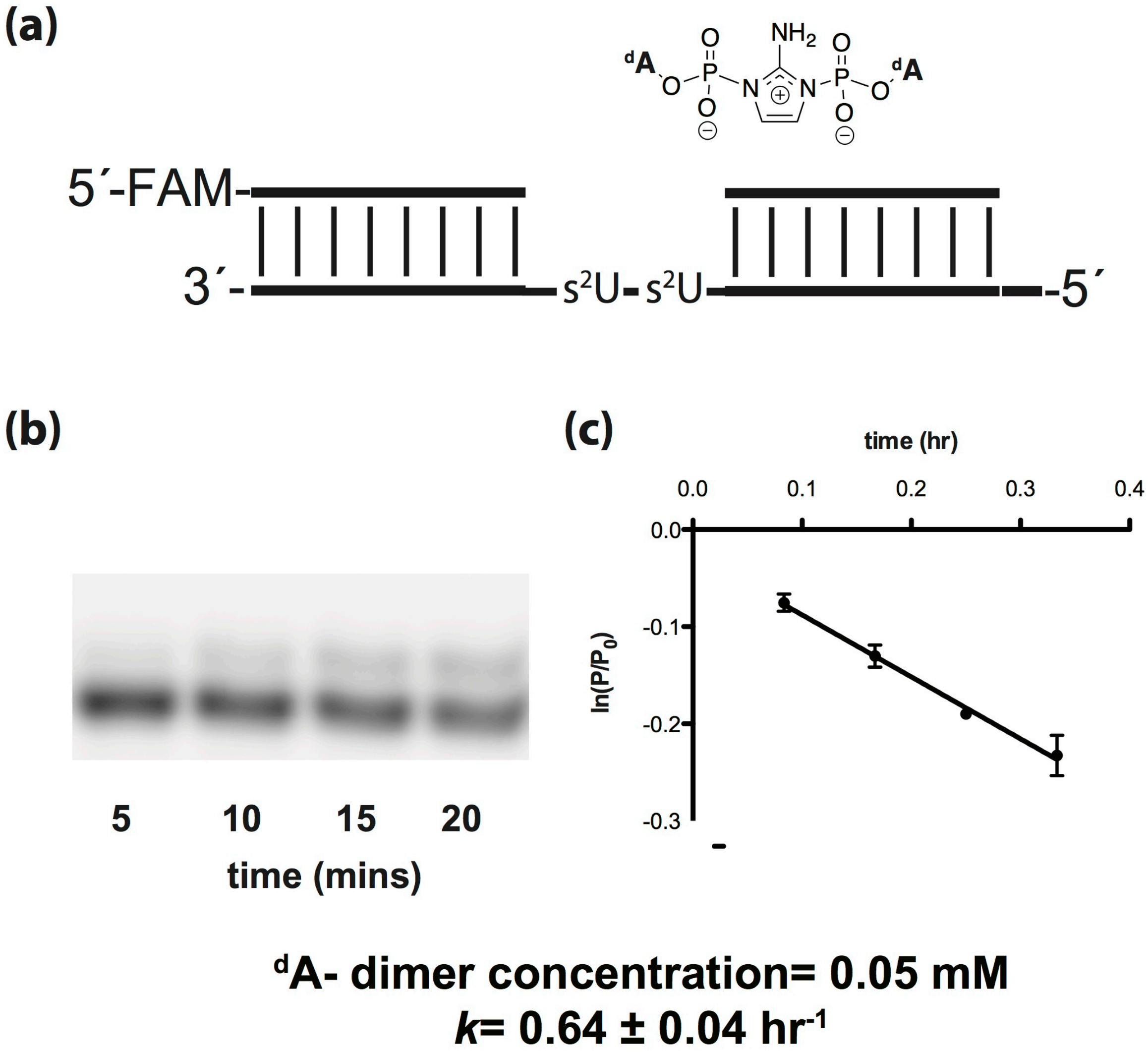


**Figure S24.** Michaelis-Menten kinetics of the addition of 2AIpdA imidazolium-bridged dimer. Primer extension reactions were carried out in triplicate using 0.05 mM of 2AIpdA imidazolium-bridged dimer, 50 mM $MgCl_2$, 200 mM $Na^+$-HEPES pH 8.0. (a) Schematic representation of a primer extension reaction. (b) Representative PAGE analysis of result. (c) Plot of $\ln(P/P_0)$as a function of time. The rate of extension was determined from linear least-squares fits of the data from three independent experiments.

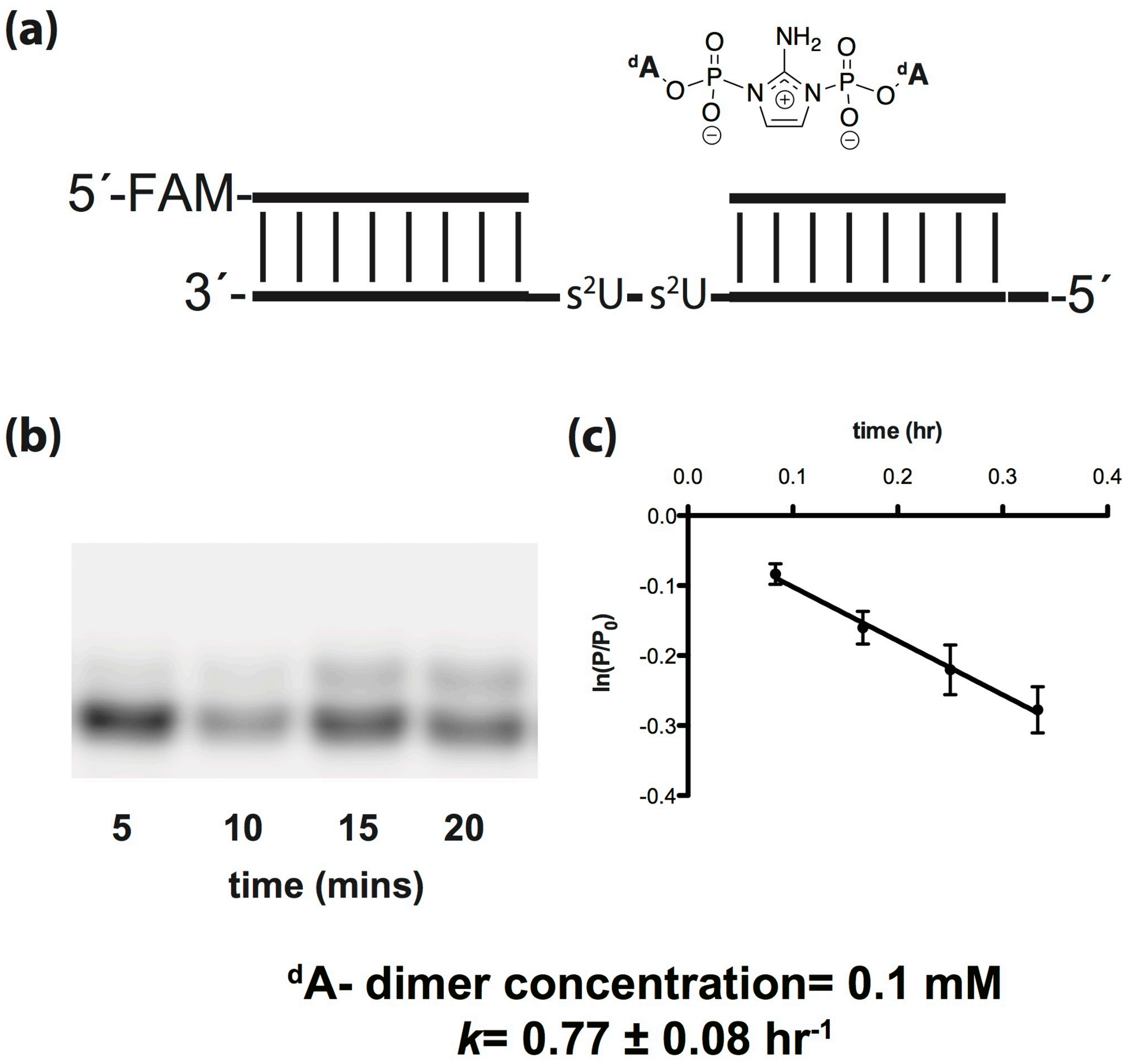


**Figure S25.** Michaelis-Menten kinetics of the addition of 2AIpdA imidazolium-bridged dimer. Primer extension reactions were carried out in triplicate using 0.1 mM of 2AIpdA imidazolium-bridged dimer, 50 mM $MgCl_2$, 200 mM $Na^+$-HEPES pH 8.0. (a) Schematic representation of a primer extension reaction. (b) Representative PAGE analysis of result. (c) Plot of $\ln(P/P_0)$as a function of time. The rate of extension was determined from linear least-squares fits of the data from three independent experiments.

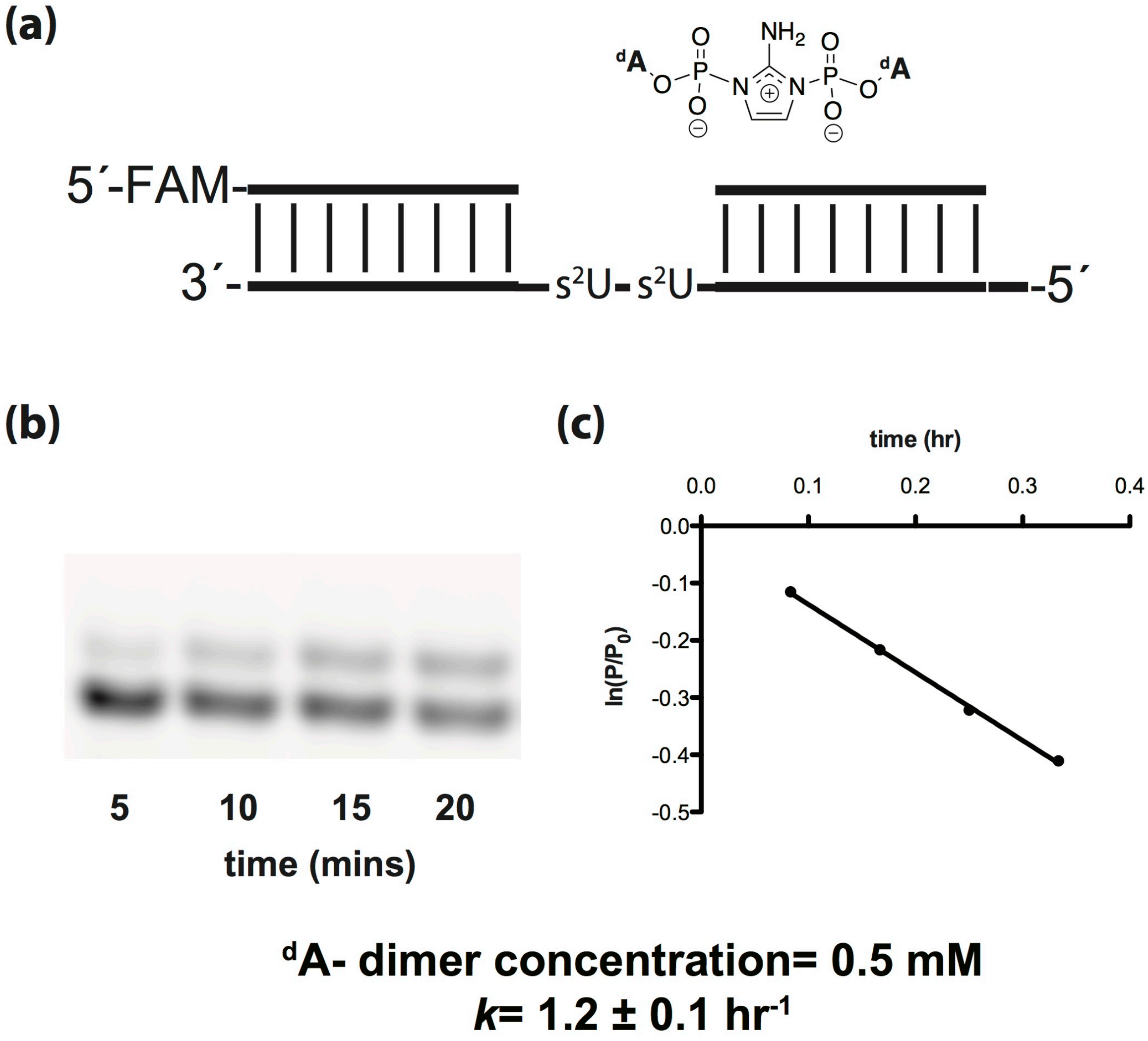


**Figure S26.** Michaelis-Menten kinetics of the addition of 2AIpdA imidazolium-bridged dimer. Primer extension reactions were carried out in triplicate using 0.5 mM of 2AIpdA imidazolium-bridged dimer, 50 mM $MgCl_2$, 200 mM $Na^+$-HEPES pH 8.0. (a) Schematic representation of a primer extension reaction. (b) Representative PAGE analysis of result. (c) Plot of $\ln(P/P_0)$as a function of time. The rate of extension was determined from linear least-squares fits of the data from three independent experiments.

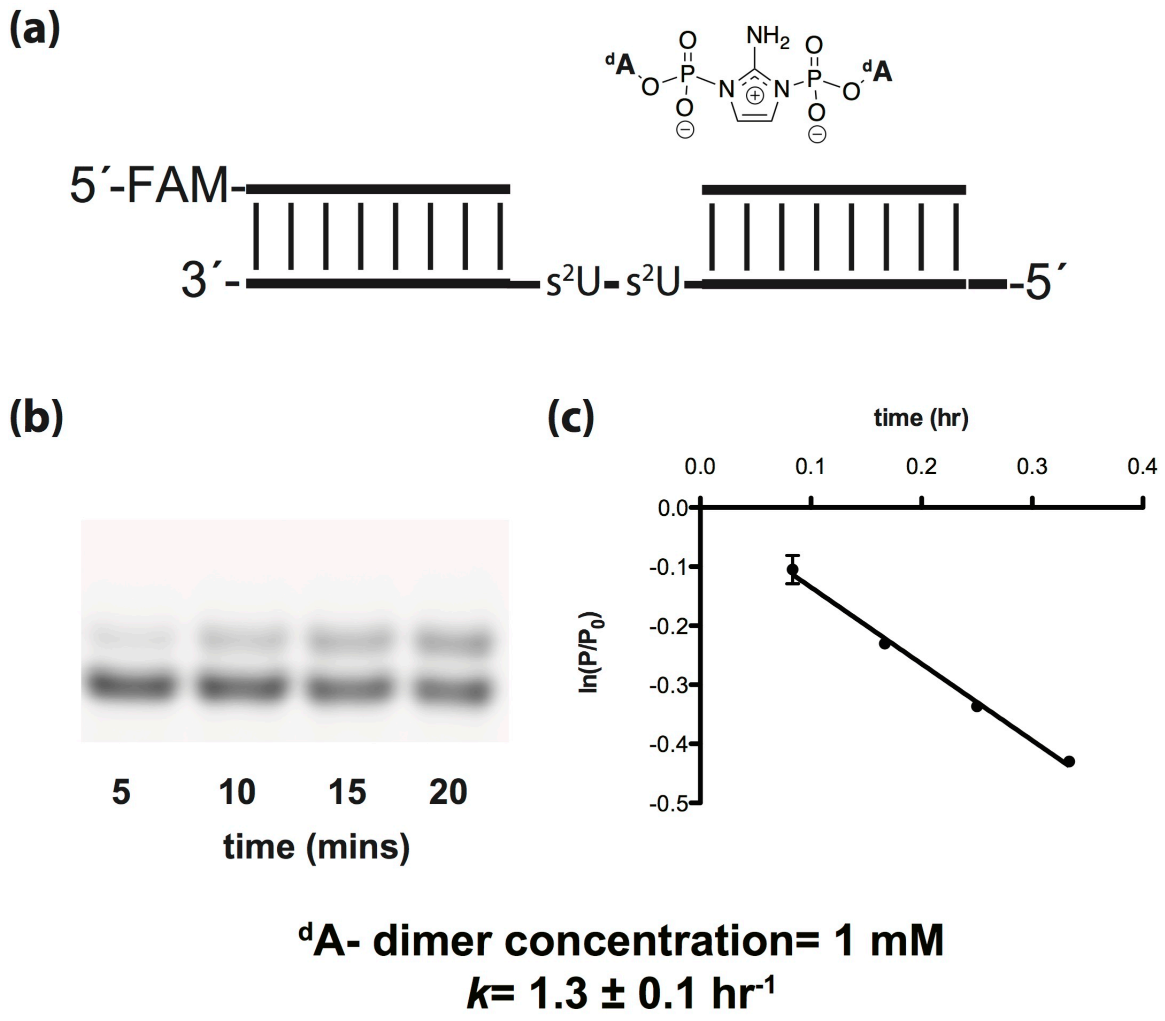


**Figure S27.** Michaelis-Menten kinetics of the addition of 2AIpdA imidazolium-bridged dimer. Primer extension reactions were carried out in triplicate using 1 mM of 2AIpdA imidazolium-bridged dimer, 50 mM $MgCl_2$, 200 mM $Na^+$-HEPES pH 8.0. (a) Schematic representation of a primer extension reaction. (b) Representative PAGE analysis of result. (c) Plot of $\ln(P/P_0)$as a function of time. The rate of extension was determined from linear least-squares fits of the data from three independent experiments.

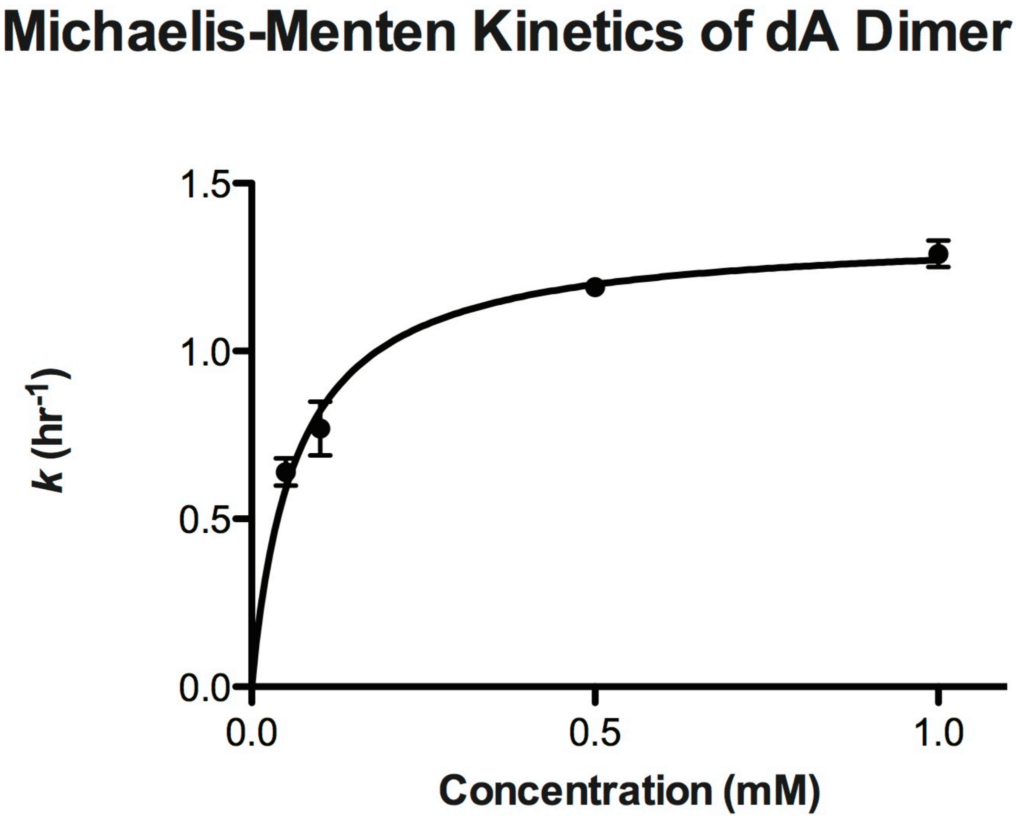


$$K_m = 0.065 \pm 0.007 \text{ mM hr}^{-1}$$
$$V_{max} = 1.4 \pm 0.1 \text{ hr}^{-1}$$

**Figure S28.** Michaelis-Menten kinetics of the addition of 2AIpdA imidazolium-bridged dimer. Primer extension reactions were carried out in triplicate using various concentrations of 2AIpdA imidazolium-bridged dimer, 50 mM $MgCl_2$, 200 mM $Na^+$-HEPES pH 8.0 and plot of $k$ ($hr^{-1}$) as a function of the concentration (mM) of 2AIpdA imidazolium-bridged dimer is shown as above. Michaelis-Menten parameters are $K_M$ : 0.065(7) mM and $V_{MAX}$ of 1.4(1) mM $hr^{-1}$.

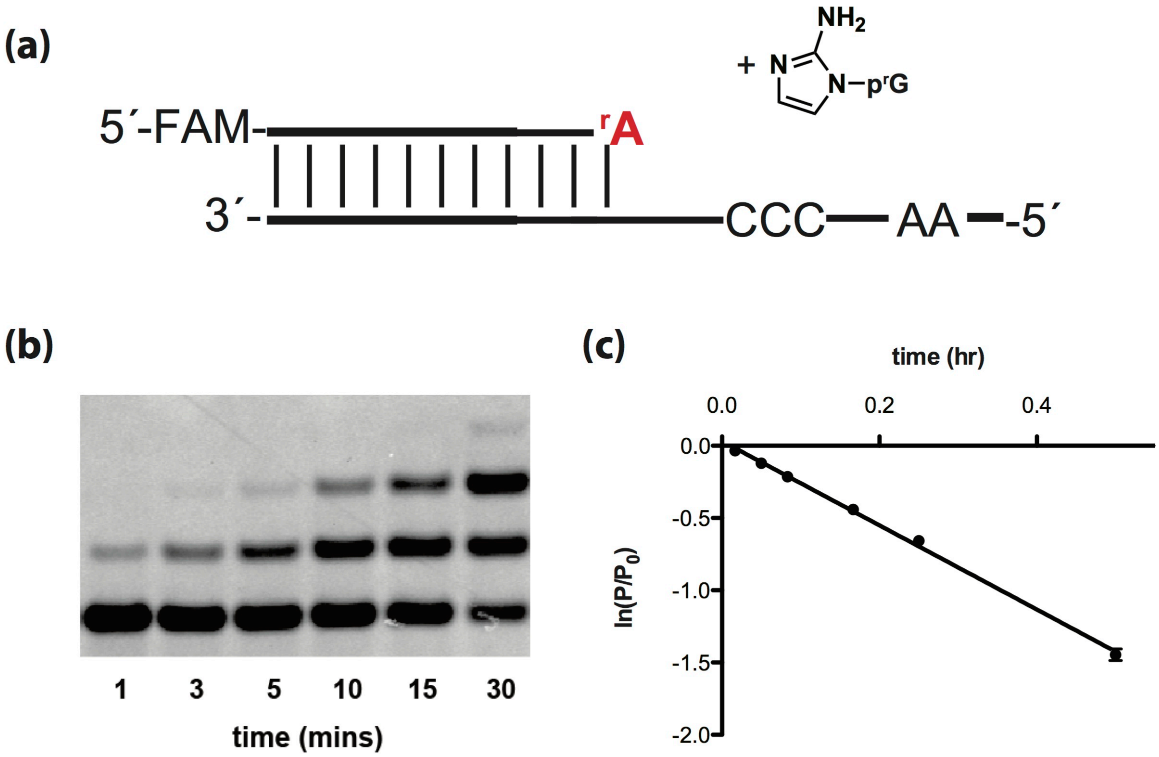


***k*= 2.8 ± 0.1 hr$^{-1}$**

**Figure S29.** Primer extension reactions were carried out in triplicate using 20 mM 2AIprG, 50 mM $MgCl_2$, 200 mM $Na^+$-HEPES pH 8.0. (a) Schematic representation of a primer extension reaction rA terminated primer. (b) Representative PAGE analysis of result. (c) Plot of ln($P/P_0$) as a function of time. The rate of extension was determined from linear least-squares fits of the data from three independent experiments.

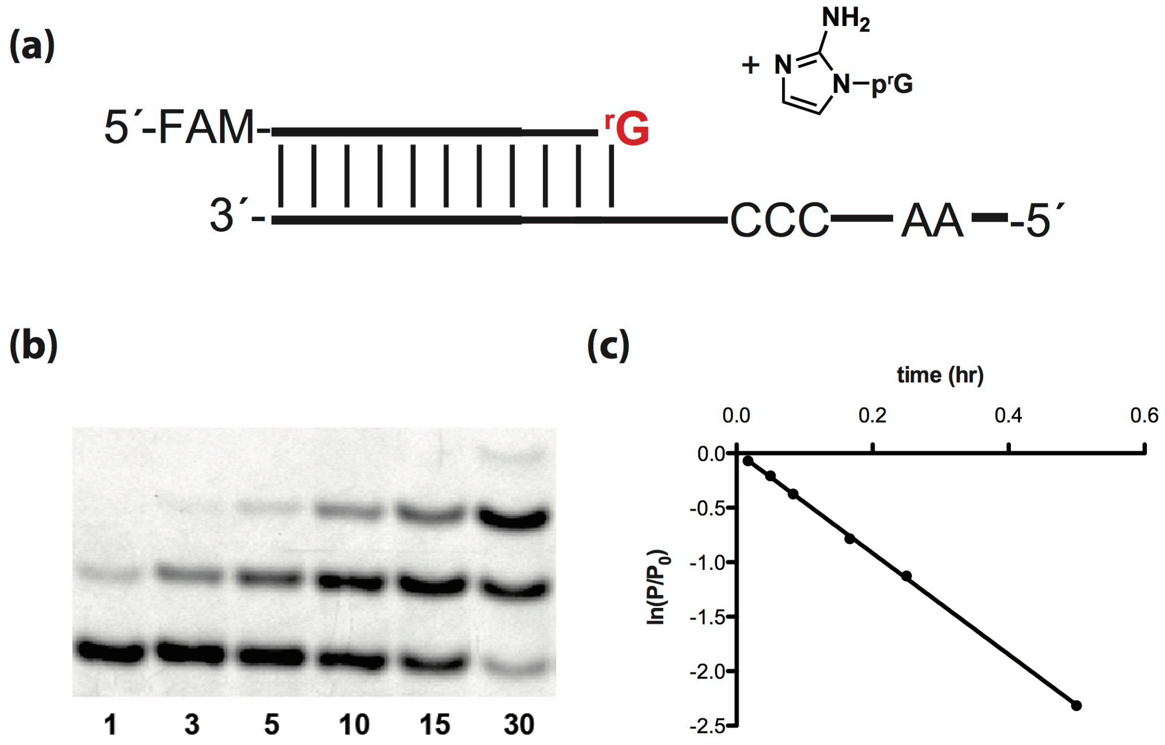


**$k$= 4.6 ± 0.1 hr$^{-1}$**

**Figure S30.** Primer extension reactions were carried out in triplicate using 20 mM 2AIprG, 50 mM $MgCl_2$, 200 mM $Na^+$-HEPES pH 8.0. (a) Schematic representation of a primer extension reaction rG terminated primer. (b) Representative PAGE analysis of result. (c) Plot of ln($P/P_0$) as a function of time. The rate of extension was determined from linear least-squares fits of the data from three independent experiments.

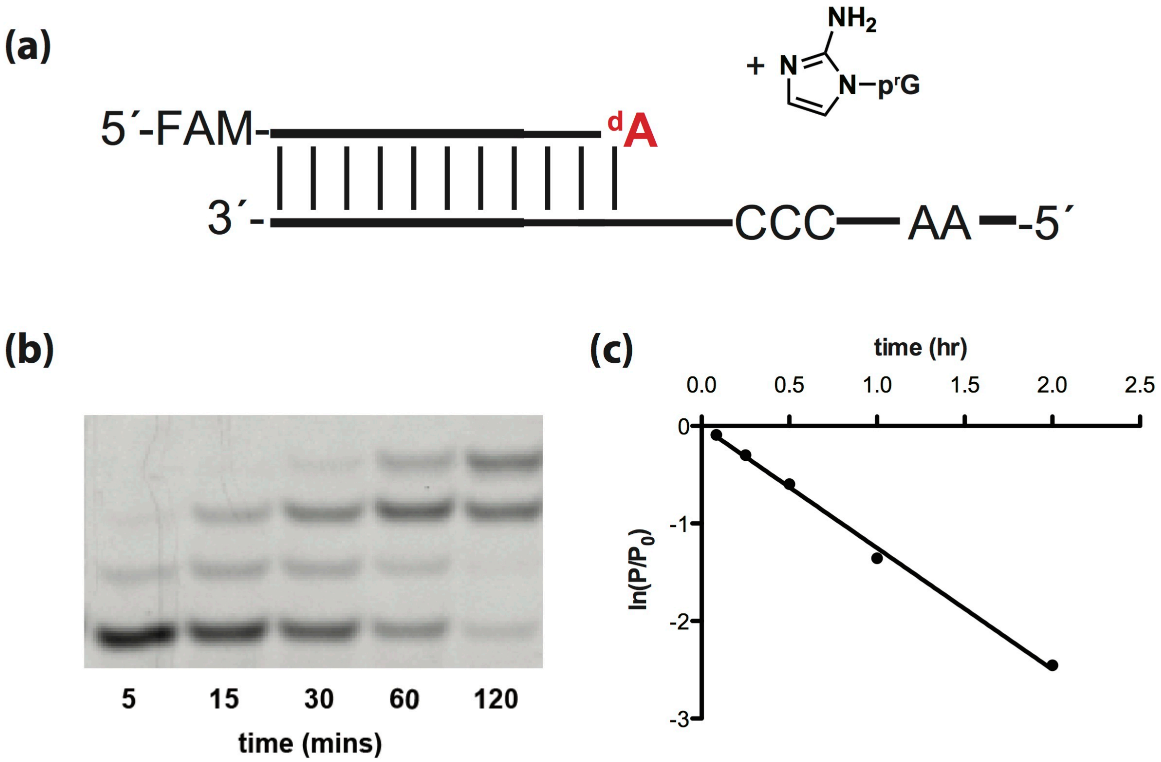


$k$= 1.3 ± 0.1 hr$^{-1}$

**Figure S31.** Primer extension reactions were carried out in triplicate using 20 mM 2AIprG, 50 mM $MgCl_2$, 200 mM $Na^+$-HEPES pH 8.0. (a) Schematic representation of a primer extension reaction dA terminated primer. (b) Representative PAGE analysis of result. (c) Plot of $\ln(P/P_0)$ as a function of time. The rate of extension was determined from linear least-squares fits of the data from three independent experiments.

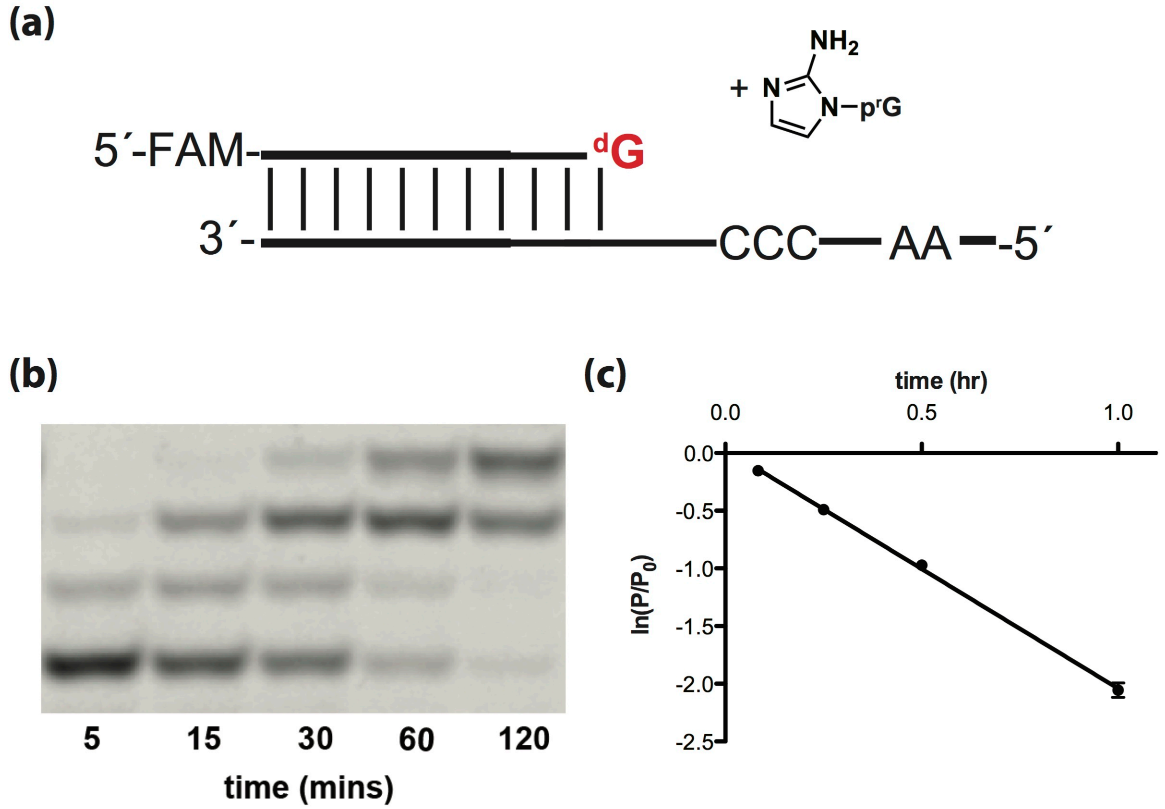


**$k$= 2.1 ± 0.1 hr$^{-1}$**

**Figure S32.** Primer extension reactions were carried out in triplicate using 20 mM 2AIprG, 50 mM $MgCl_2$, 200 mM $Na^+$-HEPES pH 8.0. (a) Schematic representation of a primer extension reaction dG terminated primer. (b) Representative PAGE analysis of result. (c) Plot of ln($P/P_0$) as a function of time. The rate of extension was determined from linear least-squares fits of the data from three independent experiments.

(a)

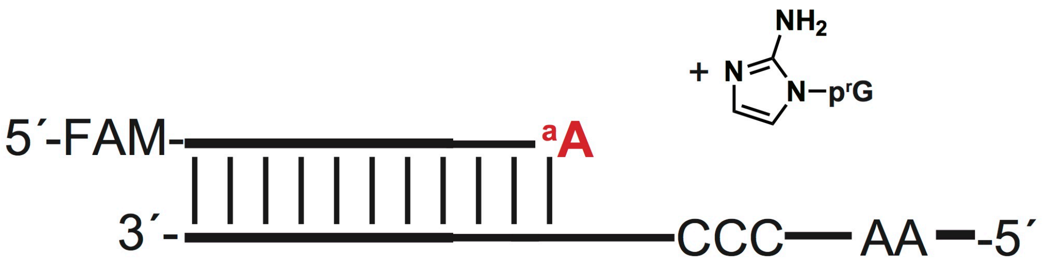


(b)

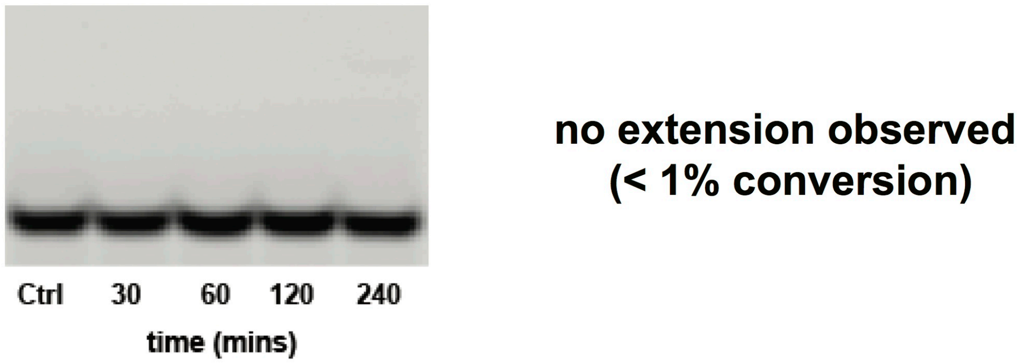


**Figure S33.** Primer extension reactions were carried out in triplicate using 20 mM 2AIprG, 50 mM $MgCl_2$, 200 mM $Na^+$-HEPES pH 8.0. (a) Schematic representation of a primer extension reaction araA terminated primer. (b) Representative PAGE analysis of result.

(a)

(b)

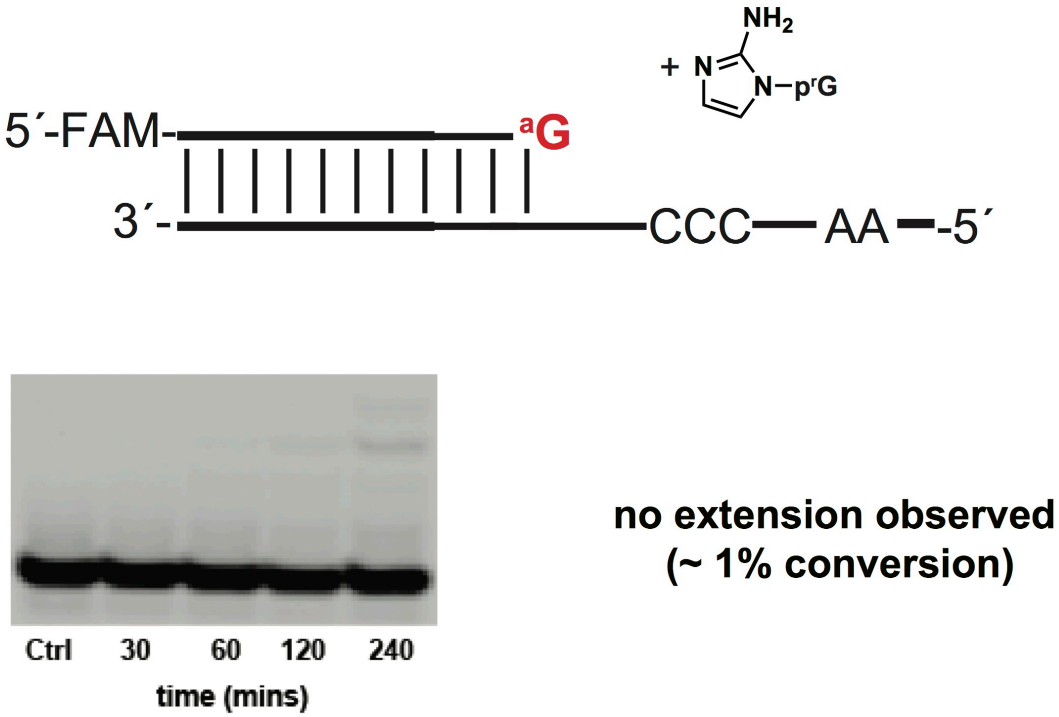


**Figure S34.** Primer extension reactions were carried out in triplicate using 20 mM 2AIprG, 50 mM $MgCl_2$, 200 mM $Na^+$-HEPES pH 8.0. (a) Schematic representation of a primer extension reaction araG terminated primer. (b) Representative PAGE analysis of result.

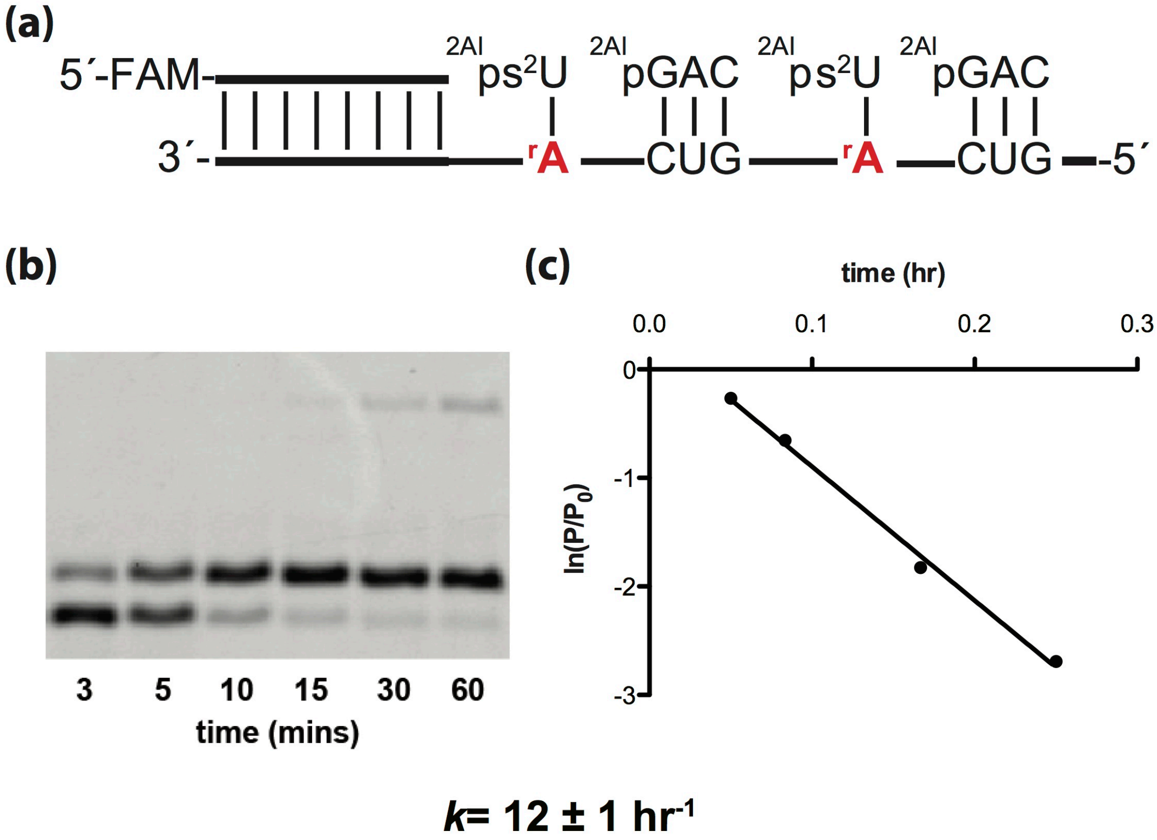


**Figure S35.** Primer extension reactions were carried out in triplicate using 20 mM 2AIps$^2$U, 0.5 mM 2AIpGAC, 50 mM $MgCl_2$, 200 mM $Na^+$-HEPES pH 8.0. (a) Schematic representation of a primer extension reaction using rA template. (b) Representative PAGE analysis of result. (c) Plot of ln($P/P_0$) as a function of time. The rate of extension was determined from linear least-squares fits of the data from three independent experiments.

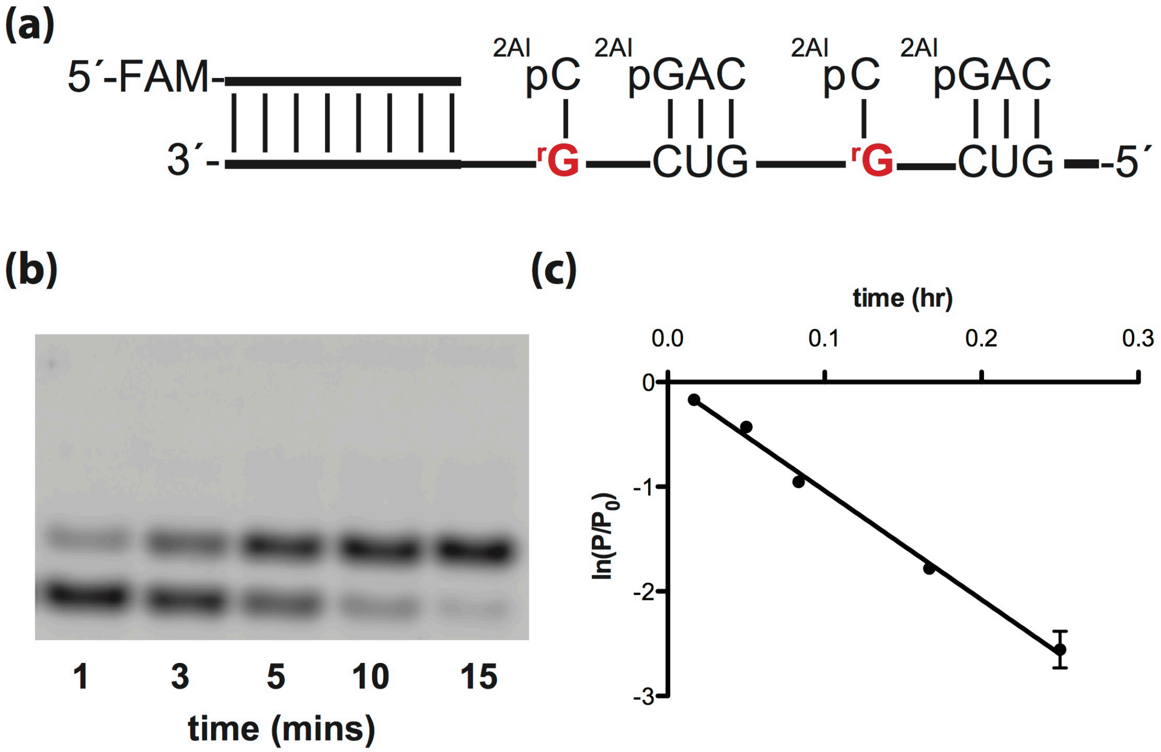


***k*= 10 ± 1 hr$^{-1}$**

**Figure S36.** Primer extension reactions were carried out in triplicate using 20 mM 2AIpC, 0.5 mM 2AIpGAC, 50 mM $MgCl_2$, 200 mM $Na^+$-HEPES pH 8.0. (a) Schematic representation of a primer extension reaction using rG template. (b) Representative PAGE analysis of result. (c) Plot of ln($P/P_0$) as a function of time. The rate of extension was determined from linear least-squares fits of the data from three independent experiments.

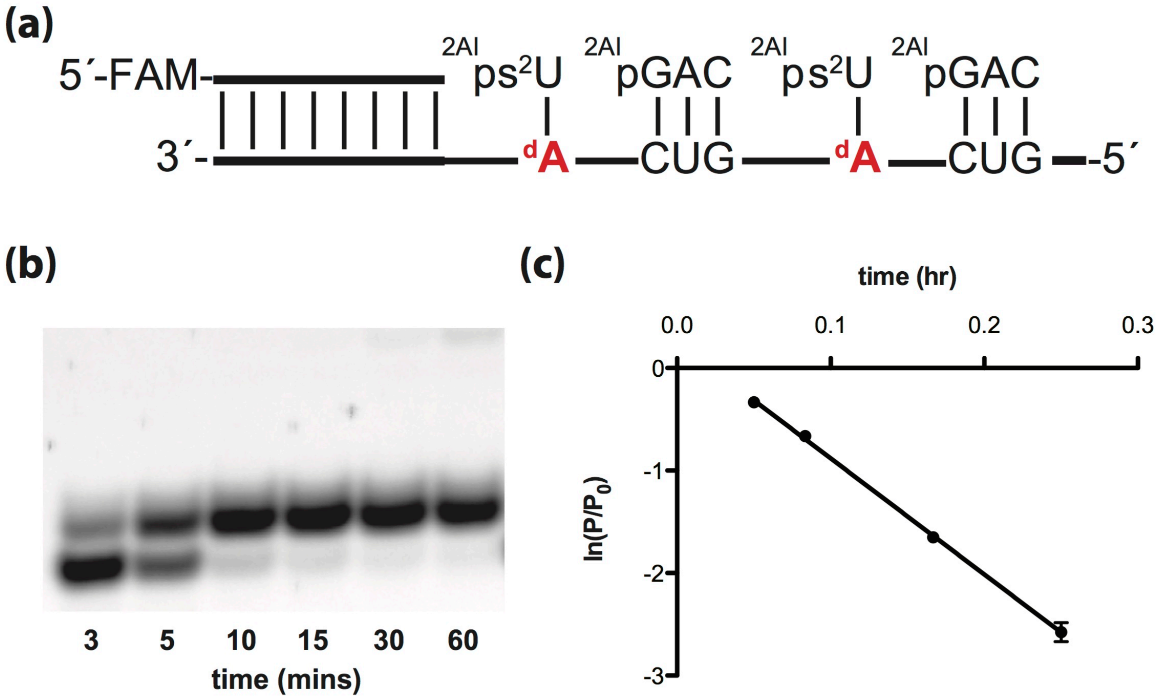


$$k = 11 \pm 1\ \text{hr}^{-1}$$

**Figure S37.** Primer extension reactions were carried out in triplicate using 20 mM 2AIps$^2$U, 0.5 mM 2AIpGAC, 50 mM $MgCl_2$, 200 mM $Na^+$-HEPES pH 8.0. (a) Schematic representation of a primer extension reaction using dA template. (b) Representative PAGE analysis of result. (c) Plot of ln($P/P_0$) as a function of time. The rate of extension was determined from linear least-squares fits of the data from three independent experiments.

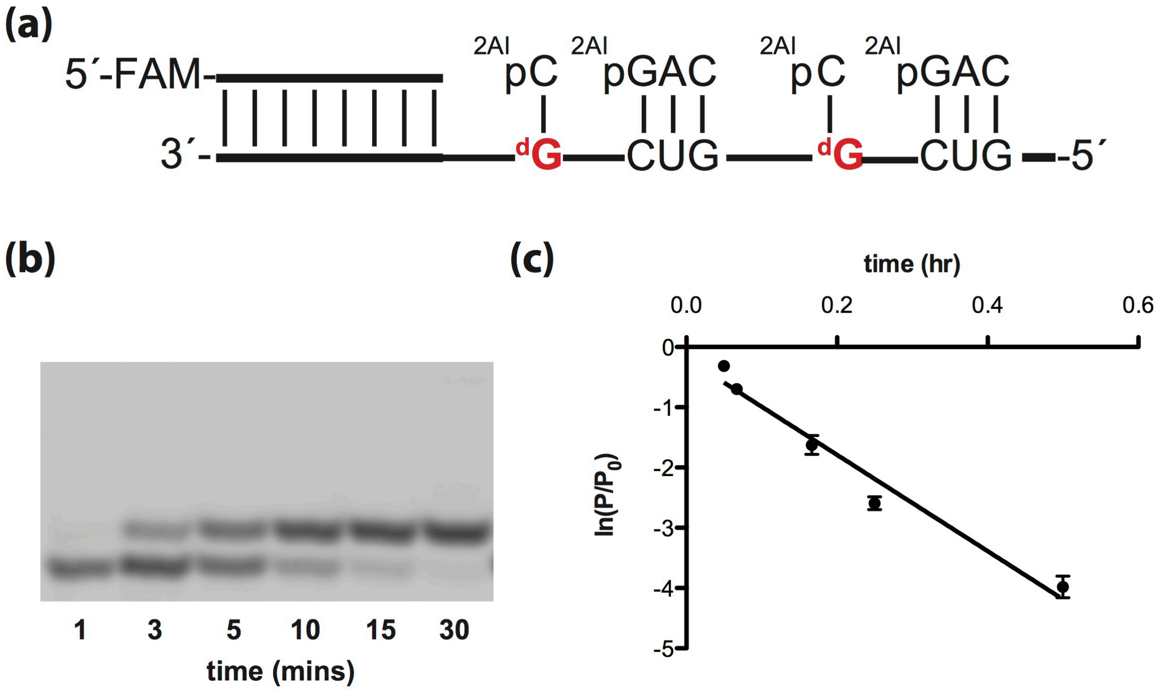


***k*= 8.0 ± 0.4 hr$^{-1}$**

**Figure S38.** Primer extension reactions were carried out in triplicate using 20 mM 2AIpC, 0.5 mM 2AIpGAC, 50 mM $MgCl_2$, 200 mM $Na^+$-HEPES pH 8.0. (a) Schematic representation of a primer extension reaction using dG template. (b) Representative PAGE analysis of result. (c) Plot of ln($P/P_0$) as a function of time. The rate of extension was determined from linear least-squares fits of the data from three independent experiments.

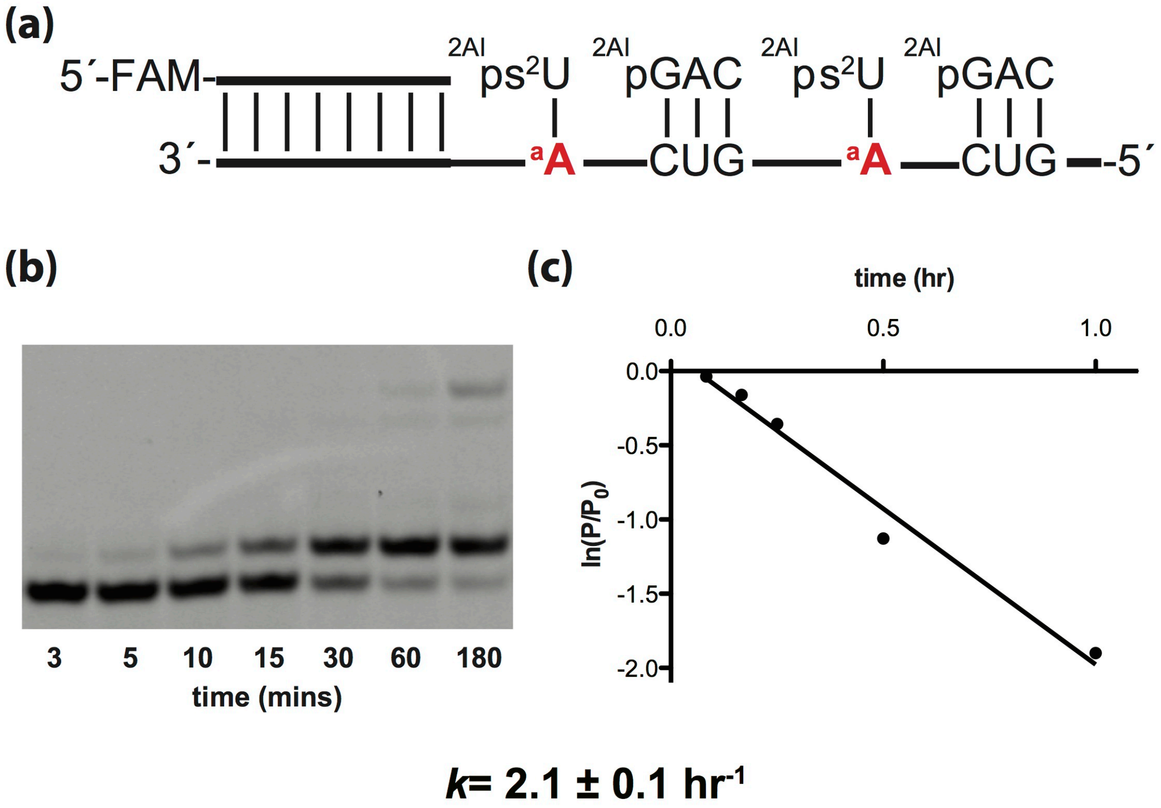


**Figure S39.** Primer extension reactions were carried out in triplicate using 20 mM 2AIps$^2$U, 0.5 mM 2AIpGAC, 50 mM $MgCl_2$, 200 mM $Na^+$-HEPES pH 8.0. (a) Schematic representation of a primer extension reaction using araA template. (b) Representative PAGE analysis of result. (c) Plot of $\ln(P/P_0)$ as a function of time. The rate of extension was determined from linear least-squares fits of the data from three independent experiments.

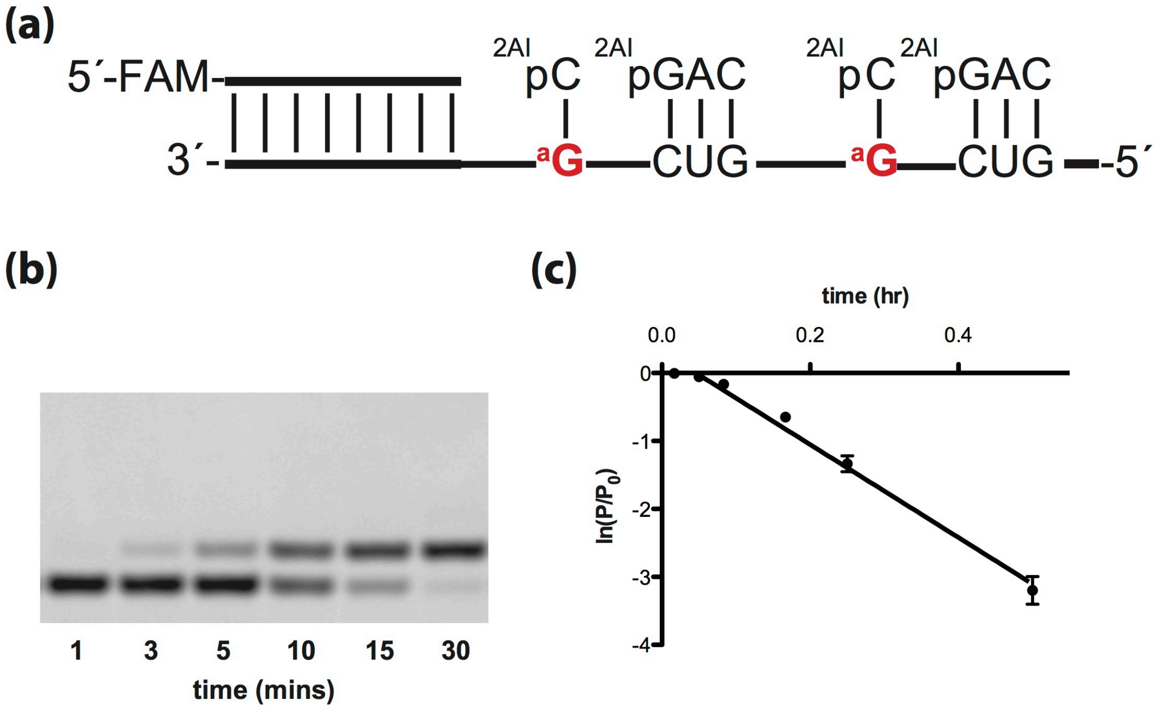


***k*= 6.8 ± 0.2 hr$^{-1}$**

**Figure S40.** Primer extension reactions were carried out in triplicate using 20 mM 2AIpC, 0.5 mM 2AIpGAC, 50 mM $MgCl_2$, 200 mM $Na^+$-HEPES pH 8.0. (a) Schematic representation of a primer extension reaction using araG template. (b) Representative PAGE analysis of result. (c) Plot of $\ln(P/P_0)$ as a function of time. The rate of extension was determined from linear least-squares fits of the data from three independent experiments.
